# Minimal lactazole scaffold for *in vitro* production of pseudo-natural thiopeptides

**DOI:** 10.1101/807206

**Authors:** Alexander A. Vinogradov, Morito Shimomura, Yuki Goto, Taro Ozaki, Shumpei Asamizu, Yoshinori Sugai, Hiroaki Suga, Hiroyasu Onaka

**Affiliations:** Department of Chemistry, Graduate School of Science, The University of Tokyo, Bunkyo-ku, Tokyo 113-0033, Japan; Department of Biotechnology, Graduate School of Agricultural and Life Sciences, The University of Tokyo, Bunkyo-ku, Tokyo 113-8657, Japan; Collaborative Research Institute for Innovative Microbiology, The University of Tokyo 1-1-1 Yayoi, Bunkyo-ku, Tokyo 113-8657, Japan

## Abstract

Lactazole A is a cryptic thiopeptide from *Streptomyces lactacystinaeus*, encoded by a compact 9.8 kb biosynthetic gene cluster. Here, we established a platform for *in vitro* biosynthesis of lactazole A, referred to as the FIT-Laz system, via a combination of the flexible *in vitro* translation (FIT) system with recombinantly produced lactazole biosynthetic enzymes. Systematic dissection of lactazole biosynthesis revealed remarkable substrate tolerance of the biosynthetic enzymes, and led to the development of the “minimal lactazole scaffold”, a construct requiring only 6 post-translational modifications for macrocyclization. Efficient assembly of such minimal thiopeptides with FIT-Laz enabled access to diverse lactazole analogs with 10 consecutive mutations, 14- to 62-membered macrocycles, and up to 18 amino acid-long tail regions. Moreover, utilizing genetic code reprogramming, we demonstrated synthesis of pseudo-natural lactazoles containing 4 non-proteinogenic amino acids. This work opens possibilities in exploring novel sequence space of pseudo-natural thiopeptides.

## Introduction

Thiopeptides are natural products defined by a six-membered nitrogenous heterocycle, usually pyridine, grafted within the backbone of a peptidic macrocycle.^1^ Multiple azole rings, dehydroamino acids, and other optional nonproteinogenic elements further contribute to the resulting structural complexity characteristic of thiopeptides. More than a hundred thiopeptides isolated to date are defined by strong antibiotic activity against Gram-positive bacteria, including methicillin resistant *Staphyrococcus aureus* (MRSA).^1–3^ For instance, thiostrepton has been used as a topical antibiotic in veterinary medicine, and LFF571, a synthetic derivative of naturally occurring GE2270A, underwent clinical trials as a treatment against *Clostridium difficile* infections.^4^

A decade ago, thiopeptides were shown to be of ribosomal origin.^5–8^ During biosynthesis, a structural gene encoding a thiopeptide precursor is transcribed and translated, and the resulting peptide undergoes post-translational modifications (**PTM**s) introduced by cognate enzymes colocalized with the structural gene in a biosynthetic gene cluster (**BGC**). Commonly, these enzymes utilize the N-terminal leader peptide (**LP**) region of the precursor as a recognition sequence, and act on the core peptide (**CP**) to introduce PTMs such as azole and dehydroalanine (**Dha**). Eventually, a pyridine synthase catalyzes formation of a six-membered heterocycle in the CP and eliminates the LP, yielding a macrocyclic thiopeptide. Thus, thiopeptides represent a group of ribosomally synthesized and post-translationally modified peptide (**RiPP**) natural products.^9^

RiPP biosynthetic logic is highly conducive to bioengineering.^10,11^ Simple nucleotide substitutions in the structural gene yield novel compounds, provided that these mutations are tolerated by the biosynthetic machinery. For BGCs encoding promiscuous enzymes, e.g. lanthipeptides and cyanobactins, this strategy can be applied to construct combinatorial libraries of natural product analogs. Recent studies demonstrated that such libraries can be screened to improve or completely reprogram antibacterial activities of the underlying RiPPs.^12–19^

In contrast, thiopeptide bioengineering proved to be significantly more challenging. Single-point mutagenesis studies^20–24^ and a few complementary reports (e.g., BGC minimization,^25^ and an incorporation of a single non-proteinogenic amino acid (**npAA**) suitable for bioconjugation)^26^ represent the bulk of the work on this topic. The challenges in thiopeptide bioengineering can be attributed to a highly cooperative yet only partially understood biosynthesis process. For many thiopeptides, the roles of individual biosynthetic enzymes are only beginning to be elucidated.^27–31^ Chemoenzymatic and semisynthetic approaches^32–38^ may circumvent the limitations imposed by biosynthetic machinery, but due to the structural complexity of thiopeptides, these strategies present a number of challenges of their own.

We previously reported isolation and characterization of lactazole A, a cryptic thiopeptide from *Streptomyces lactacystinaeus* (Fig. 1a).^39^ It is biosynthesized from a compact 9.8 kb *laz* BGC encoding just five enzymes essential for the macrocycle formation (Fig. 1b). Lactazole A has a low Cys/Ser/Thr content, a 32-membered macrocycle, and bears an unmodified amino acid in position 2 (Trp2), all of which are unusual features among thiopeptides (Fig. 1c).^40^ Moreover, lactazole A shows no antibacterial activity, and its primary biological function remains unknown. Recent bioinformatic studies indicated that the lactazole-like thiopeptides, characterized by an unmodified amino acid in position 2, comprise close to half of all predicted thiopeptides (251 out of 508 annotated BGCs), and yet the prototypical *laz* BGC remains the only characterized member of this family to date.^41^ Overall, lactazole-like thiopeptides remain a rather enigmatic family of natural products, as close to nothing is known about their function, structural diversity, and biosynthesis.

**Figure 1.**
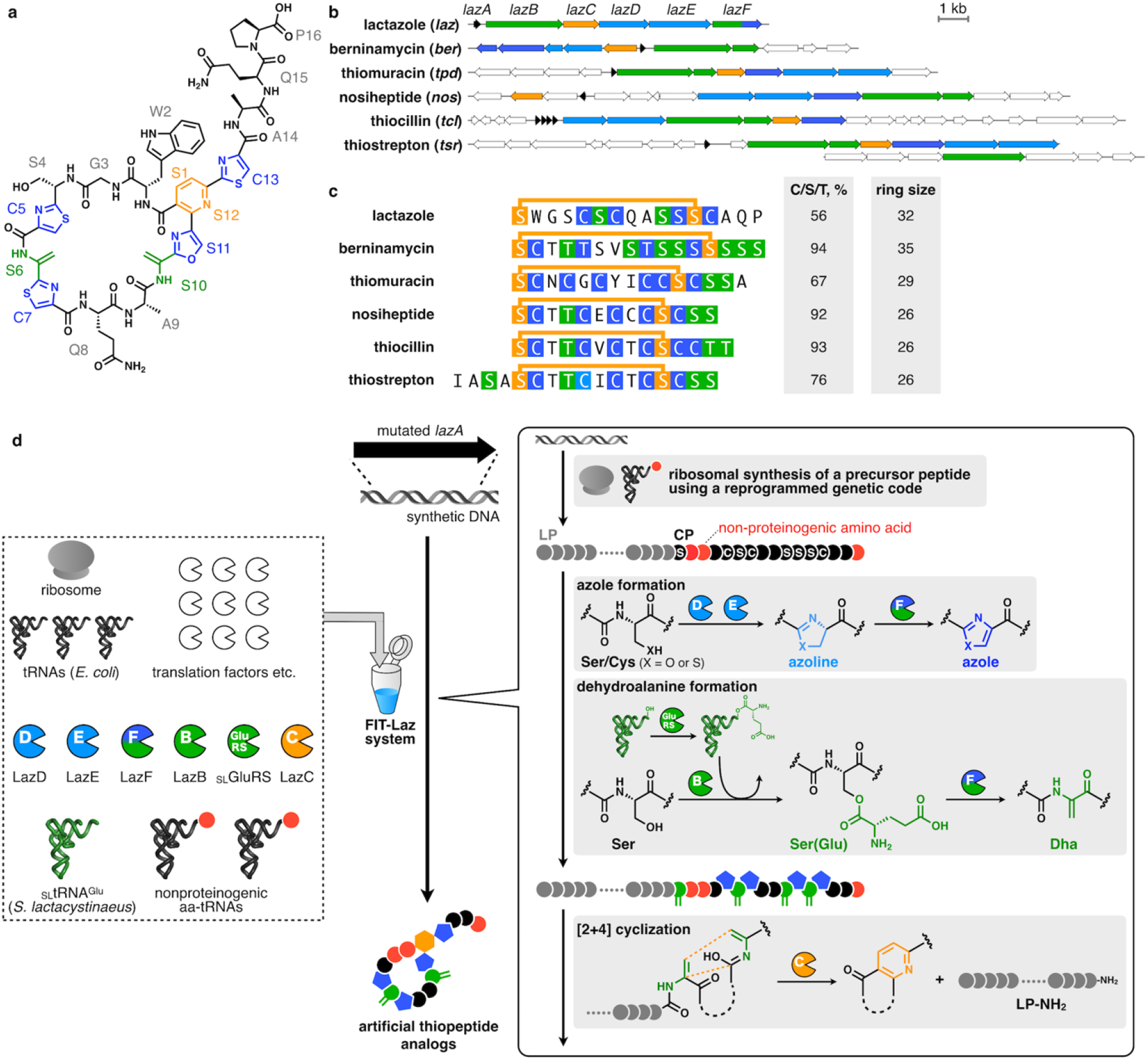
Lactazole A and its biosynthesis with the FIT-Laz system. **(a)** Chemical structure of lactazole A. **(b)** Comparison of *laz* BGC to other prototypical thiopeptide BGCs. Homologs of *laz* genes are color-coded. Genes encoding enzymes responsible for the installation of azolines, azoles, dehydroalanine, and pyridine are shown in light blue, blue, green, and orange, respectively. Precursor peptide structural genes are shown in black, and ancillary genes absent from *laz* BGC are in white. **(c)** Comparison of primary sequences for thiopeptides from panel (b), with the same PTM color coding. The comparison reveals an unusual macrocycle size, low C/S/T content and the absence of azole modification in position 2 as unique features of lactazole. **(d)** Summary of the FIT-Laz system and the roles of individual enzymes during lactazole biosynthesis. In FIT-Laz, synthetic DNA templates encoding LazA or its mutants are *in vitro* transcribed and translated to generate precursor peptides, which undergo a cascade of PTMs introduced by lactazole biosynthetic enzymes to yield lactazole A or its artificial analogs.

Intrigued by the uniqueness of *laz* BGC, we set out to reconstitute *in vitro* biosynthesis of lactazole A. We sought to establish rapid and reliable access to lactazole A and its analogs in order to evaluate the applicability of *laz* BGC for bioengineering, and to pave the way for future characterization of enzymes and BGCs from the lactazole family. To this end, we report construction of the FIT-Laz system, a combination of flexible *in vitro* translation (**FIT**)^42^ with PTM enzymes from *laz* BGC, as a platform for facile *in vitro* synthesis of lactazole-like thiopeptides (Fig. 1d). Taking advantage of the FIT-Laz system, we explored the substrate plasticity of *laz* BGC, and found that *in vitro* lactazole biosynthesis is remarkably tolerant to mutation, insertion and deletion of multiple amino acids, including npAAs. A systematic dissection of the pathway led to the identification of the “minimal lactazole scaffold”, a CP with only 5 amino acids indispensable for the macrocyclization process. Our work opens a possibility to tap into an unexplored sequence space of pseudo-natural thiopeptides, and use them as molecular scaffolds in drug lead discovery against protein targets of choice.

## Results

### *In vitro* reconstitution of lactazole biosynthesis

We began with recombinant production of Laz enzymes in *Escherichia coli* BL21(DE3). The five enzymes (LazB, LazC, LazD, LazE, and LazF) were expressed and purified as soluble His-tagged proteins (**Fig. S1**). The FIT system was used to establish access to the precursor peptide (LazA;^43^ Fig. 2a). Linear DNA template encoding LazA was assembled by PCR and incubated with the *in vitro* reconstituted translation machinery from *E. coli* supplemented with T7 RNA polymerase. This scheme for precursor peptide production parallels the previously established FIT-PatD and FIT-GS systems, used for the synthesis of azoline-containing peptides^44^ and goadsporin analogs^45^, respectively.

**Figure 2.**
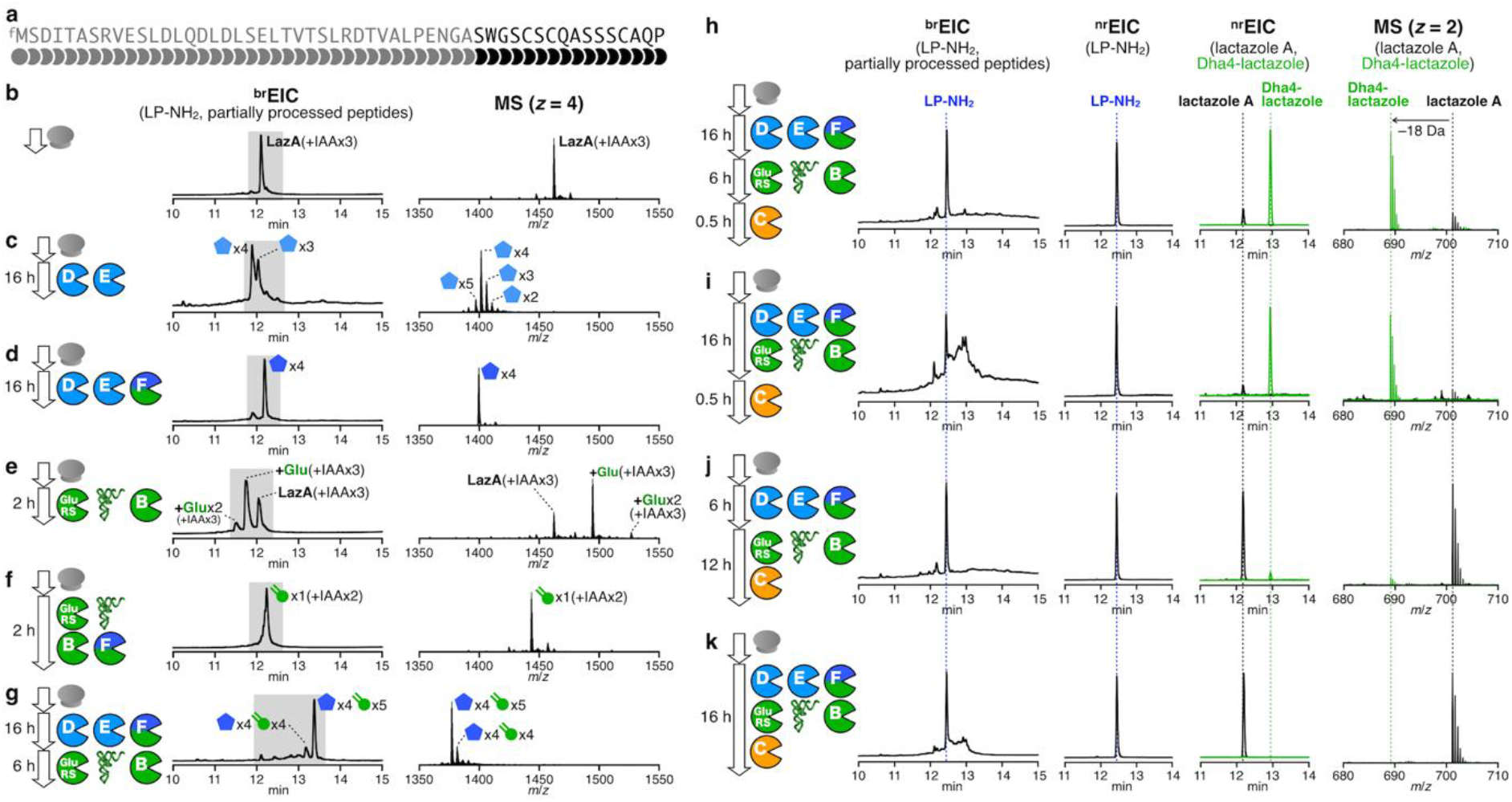
Reconstitution of *in vitro* lactazole A biosynthesis. (**a**) Primary amino acid sequence of LazA precursor peptide. **(b)** – **(g)** Reconstitution of azole and Dha formation in FIT-Laz. LazA precursor peptide produced with the FIT system was treated with a combination of Laz enzymes as indicated in each panel and the reaction outcomes were analyzed by LC-MS. Displayed are ^br^EIC LC-MS chromatograms and composite mass spectra integrated over a time period shaded in the corresponding chromatograms. See S.I 2.6 and 2.10 for details on reaction conditions and the explanation of ^br^EIC chromatograms. **(h)** – **(k)** Reconstitution of lactazole A biosynthesis in FIT-Laz. Displayed are LC-MS chromatograms (left to right: ^br^EIC; ^nr^EIC at *m/z* 1026.77 for LP-NH_2_ generated during the final macrocyclization step; overlaid ^nr^EICs at *m/z* 701.20 for lactazole A shown in black, and at *m/z* 692.20 for Dha4-lactazole in green) and overlaid mass spectra for the lactazole A and Dha4-lactazole. These results demonstrate that the order of enzyme addition is critical to the success of lactazole A biosynthesis.

With all components in hand, we turned to reconstitution of lactazole biosynthesis. Maturation of goadsporin, a distantly related linear azole-containing RiPP, is initiated with the formation of azoles, while Dha installation is dependent on it,^46^ and biosynthesis of thiomuracin also follows a similar modification order.^47^ Based on these results, we hypothesized that azole formation is the starting point in lactazole biosynthesis, and therefore attempted to reconstitute the activity of LazDEF (LazD, LazE and LazF) first. LazDE is a split YcaO cyclodehydratase^40,48^ characteristic of thiopeptide BGCs: LazD is predicted to bear a RiPPs recognition element, necessary for LP binding, ^49^ and LazE contains an ATP-binding domain,^50^ utilized for ATP-dependent cyclodehydration of Cys/Ser residues in the CP (Fig. 1d). LazF is an unusual bifunctional protein that features a fusion between an FMN-dependent dehydrogenase which oxidizes azolines installed by LazDE to azoles,^51–53^ and a glutamate elimination domain, tentatively participating in the formation of Dha (see below). After LazA precursor peptide expressed with the FIT system (Fig. 2b) was incubated with LazDE, the mixture was treated with iodoacetamide (**IAA**) and analyzed by LC-MS. The resulting broad-range extracted ion chromatogram (^**br**^**EIC**; see **S.I. 2.10** and **Fig. S2–S5** for detailed description of ^br^EIC) indicated that the LazDE reaction yielded a mixture of two, three, four and five dehydrations (Fig. 2c). Either LazD or LazE alone had no activity (**Fig. S6a and b**). In contrast, incubating LazA with LazDEF afforded a single product containing 4 azoles (Fig. 2d). No alkylation occurred on this peptide by IAA suggesting that all Cys residues were cyclized, and MS/MS analysis of this product supported the native azole pattern, *i.e.* three thiazoles in positions 5, 7, 13 and one oxazole in position 11 (**Fig. S7**).

Next, we attempted to reconstitute the Dha-forming activity (Fig. 1d). LazBF is a split dehydratase, widely conserved in thiopeptide BGCs (Fig. 1b).^54,55^ These proteins are homologous to class I lanthipeptide dehydratases, which utilize Glu-tRNA^Glu^ to glutamylate Ser or Thr residues in the CP of a substrate (using the glutamylation domain),^46^ and then catalyze elimination of the glutamate to yield Dha (using the elimination domain). In *laz* BGC, LazB is annotated as a glutamylation domain, and the N-terminal part of LazF is an elimination domain. Even though we assumed that azole formation precedes Dha synthesis, we first attempted to test LazB activity on the unmodified LazA. Surprisingly, LazB glutamylated LazA once when incubated in the presence of synthetic tRNA^Glu^ originating from *S. lactacystinaeus* and glutamyl-tRNA synthetase (**GluRS**) from *S. lividans* (Fig. 2e), while reactions lacking any one component led to no modification (**Fig. S6f-h**). These results indicated that like homologous enzymes,^56,57^ LazB utilizes Glu-charged tRNA^Glu^ and catalyzes glutamylation of LazA. Because *E. coli* tRNA^Glu^ and GluRS present in the translation mixture were not accepted, LazB appears to be specific for the *Streptomyces* tRNA^Glu^ and GluRS. The complete dehydratase activity was reconstituted with the addition of LazF to the mixture, in which case the reaction yielded a singly dehydrated product (Fig. 2f). Extending the reaction time led to sluggish second and third dehydrations (**Fig. S6i**). These results indicated that LazBF can catalyze formation of some but not all Dha in LazA independent of azole formation.

We next studied whether LazBF forms remaining Dha in an azole-dependent fashion. To this end, we incubated the LazDEF-treated LazA, bearing 4 azoles, with LazBF, tRNA^Glu^ and GluRS. This reaction led to a major product 90 Da lighter than the 4-azole LazA, consistent with the formation of 5 Dha, suggesting that all available Ser in the CP were dehydrated (Fig. 2g). Coincubation of LazA with LazBDEF, tRNA^Glu^ and GluRS resulted in the formation of a complex mixture with the same major product (**Fig. S6j**).

Finally, we tested reconstitution of the entire biosynthetic pathway by adding LazC to the reaction. The putative pyridine synthase LazC is weakly homologous to TclM and TbtD, two well-studied enzymes catalyzing analogous reactions.^38,58^ Both enzymes are believed to initiate a [4+2]-cycloaddition reaction leading to the formation of a macrocyclic product bearing dihydropyridine, which is further aromatized by eliminating LP as a C-terminal amide (**LP-NH_2_**) to give rise to a pyridine ring. Accordingly, incubation of the aforementioned LazA bearing 4 azole/5 Dha with LazC afforded LP-NH_2_ accompanied by a thiopeptide 18 Da lighter than expected, indicating that Ser4, unmodified in lactazole A, was dehydrated (Dha4-lactazole A) (Fig. 2h and i). MS/MS analysis of this product confirmed its structure (**Fig. S11**). Lactazole A was a minor product under these reaction conditions. Changing the order of the enzyme addition (LazDEF followed by LazBC, tRNA^Glu^ and GluRS) suppressed the formation of the overdehydrated product, but still, a mixture of thiopeptides formed (Fig. 2j). In contrast to the stepwise reactions, coincubation of LazA with the full enzyme set (LazBCDEF, tRNA^Glu^ and GluRS) resulted in the formation of lactazole A and LP-NH_2_ with only a trace amount of Dha4-lactazole A (Fig. 2k). LC-MS analysis of the *in vitro* synthesized lactazole A showed that its molecular weight and HPLC retention time were identical to the authentic *in vivo* synthetized standard (**Fig. S9**). Additionally, both samples had matching, annotatable CID MS/MS spectra (**Fig. S8 and S10**), indicating that the one-pot reaction yielded the authentic thiopeptide.

In summary, here we demonstrated that the translation product of *lazA* accessed with the FIT system can be treated with the full set of Laz enzymes to yield lactazole A. We refer to this series of transformations as the FIT-Laz system.

### Analysis of substrate tolerance of Laz enzymes

To understand the overall substrate plasticity of *laz* BGC, we next investigated whether the FIT-Laz system can produce lactazole analogs. We commenced with Ala-scanning mutagenesis and prepared 14 single-point Ala mutants in the CP region of LazA. The precursor peptides were expressed and modified with the FIT-Laz system, and the reaction outcomes were analyzed by LC-MS as above (Fig. 3a). Only 4 Ala mutants, S1A, S10A, S11A, and S12A, abolished formation of thiopeptides, whereas other constructs led to the formation of corresponding lactazole analogs and LP-NH_2_. During maturation, Ser1 and Ser12 are converted to Dha and are then utilized by LazC for pyridine formation/macrocyclization. Moreover, pyridine synthases are known to recognize the modification pattern around the 4π component,^38^ which is consistent with the abrogation of biosynthesis in S10A and S11A mutants. On the other hand, C13A mutant was converted to a thiopeptide without modifications in the tail region, suggesting that the Dha10-oxazole11-Dha12 moiety is the minimal recognition motif around the 4π component for the LazC-catalyzed macrocyclization. Significant accumulation of linear side-products and partially processed peptides for W2A and G3A mutants indicates that these amino acids are also important for smooth lactazole biosynthesis. Ala mutants in positions 4–8, 15 and 16, including those disrupting azole and Dha installation, were tolerated, albeit in some cases a mixture of thiopeptides formed (**Fig. S12 and S13**).

**Figure 3.**
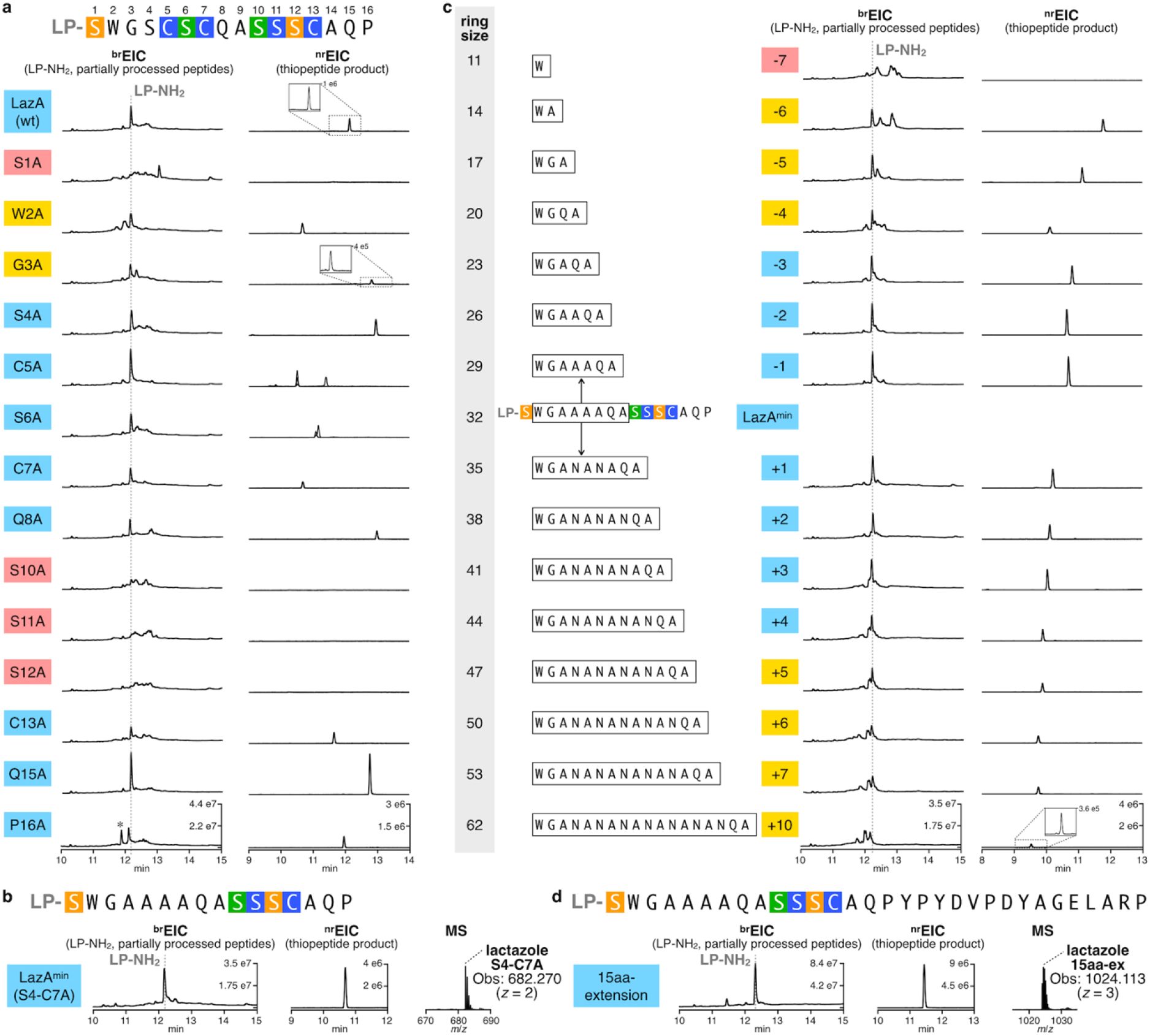
Substrate scope of the FIT-Laz system. **(a)** Ala scanning of the LazA CP. Single-point Ala mutants of LazA were treated with the full enzyme set and the outcomes were analyzed by LC-MS. Displayed are LC-MS chromatograms (^br^EIC chromatograms on the left showing partially processed linear peptides and LP-NH_2_ after enzymatic treatment, and ^nr^EIC chromatograms on the right for expected thiopeptides generated at *m/z* 0.10 tolerance window). For mutants highlighted in light blue biosynthesis proceeded efficiently; yellow highlighting indicates inefficient thiopeptide formation accompanied by the accumulation of linear intermediates and side-products; red – mutants that failed to yield a detectable thiopeptide. Peaks denoted with an asterisk (*) indicate translation side-products. Mutants C5A and S6A gave 4 and 2 thiopeptides, respectively, annotations of which can be found in **Fig. S12 and S13**. Y-axes are scaled between samples for each chromatogram type. **(b)** LC-MS chromatograms as in (a) for the enzymatic processing of LazA^min^ on the left with a zoomed-in mass spectrum of the produced thiopeptide on the right. **(c)** LC-MS chromatograms as in (a) for ring expansion and contraction study of LazA^min^. **(d)** LC-MS chromatograms and mass spectrum as in (b) for a LazA^min^ variant containing a 15-amino acid extension in the tail region. Collectively, these data point to the remarkable substrate tolerance of Laz enzymes.

Intrigued by these results, we examined whether non-essential modifications inside the macrocycle (Ser4–Cys7) can be removed altogether. Indeed, a tetra-Ala mutant, LazA S4-C7A, was converted to a thiopeptide containing just 2 azoles and 1 Dha upon treatment with the full enzyme set (Fig. 3b). A pentamutant LazA S4-C7A, C13A also afforded a thiopeptide, but at a much lower overall efficiency, as a number of partially processed linear peptides accumulated after overnight treatment (**Fig. S14a**). Based on these results, we concluded that the five residues undergoing PTM in LazA S4-C7A (Ser1, Ser10, Ser11, Ser12 and Cys13) are essential for efficient maturation. We termed the resulting thiopeptide as the minimal lactazole scaffold, and the corresponding precursor peptide as the minimal lactazole precursor (**LazA^min^**). Because *in vitro* biosynthesis of the minimal lactazole proceeded as efficiently as the wild type, we decided to investigate enzymatic processing of LazA^min^ and its potential for bioengineering applications in more detail.

In the next series of experiments, we examined the tolerance of Laz enzymes to the presence of charged amino acids in the CP, and performed Lys-and Glu-scanning of LazA^min^ CP. Charged amino acids are rarely found in CPs of thiopeptides,^39–41^ and RiPP enzymes from other classes are also known to disfavor charged amino acids in general, especially negatively charged Asp and Glu close to the modification site. We prepared 11 single-point Lys mutants and 11 single-point Glu mutants in the non-essential positions of LazA^min^ and analyzed their processing as above (**Fig. S15**). The results of Lys-scanning revealed that a positively charged amino acid is well tolerated in 9 out of 11 positions, whereas W2K and A14K mutants suffered from inefficient processing. Glu was less accepted than Lys overall. In addition to inefficient processing of W2E and A14E, mutants Q8E, A9E and P16E also resulted in little to no thiopeptide formation. These data suggest that in addition to the five previously identified amino acids, Trp2 and Ala14 also play an important role in LazA^min^ maturation.

Next, we sought to establish the minimal and maximum macrocycle sizes accessible with FIT-Laz. All known thiopeptides range between 26-(thiocillin, thiostrepton and nosiheptide) and 35-membered macrocycles (berninamycin).^59^ Additionally, previously reported bioengineering of thiocillin BGC led to 23-membered artificial variants.^60^ Here, we prepared amino acid insertion and deletion variants of LazA^min^, and, as before, expressed and modified them with the FIT-Laz system. The results of LC-MS analysis are summarized in Fig. 3c. Deletion of up to 3 amino acids between Ser1 and Ser10 was well tolerated, and led to the efficient formation of 29-to 23-membered thiopeptides. The 4–6 amino acid deletion mutants were also competent substrates and yielded 20-to 14-membered macrocycles, but at relatively low overall efficiencies, as a number of partially processed linear peptides accumulated. Formation of an 11-membered thiopeptide (deletion of 7 residues) was not observed. Thus, it appears that 14-membered thiopeptides are the smallest accessible with Laz enzymes, 9 atoms smaller than the previously smallest thiocillin variants.^60^ In contrast, no upper limit on the ring size could be placed. All tested substrates were accepted by the enzymes: the largest synthesized product bore a 62-membered macrocycle, which corresponds to a 10 amino acid insertion. The overall processing efficiency decreased linearly with increasing the cycle size; where LazA^min^ itself was efficiently converted to a macrocycle and LP-NH_2_, substrates with multiple amino acid insertions had substantial accumulation of linear intermediates and side-products. A recent study demonstrated that TbtD, a LazC homologue from thiomuracin biosynthesis, could perform an *inter*molecular [4+2]-cycloaddition, whereas TclM, an enzyme from thiocillin BGC, could not,^61^ suggesting that LazC might function similarly to TbtD. While certainly not intermolecular, the enzyme catalyzed formation of remarkably large macrocycles. Sequence extension outside of the macrocycle was also easily achievable, as 3 LazA^min^ variants with the C-terminal tail extensions of up to 15 amino acids were efficiently converted to thiopeptides (Fig. **3d and S14b–d**).

Encouraged by these results, we examined whether FIT-Laz can accommodate sequence randomization inside the macrocycle. We prepared 10 LazA^min^ variants containing 10 consecutively randomized amino acids each, which corresponds to the simultaneous insertion of 3 amino acids and mutation of residues 3–9 in LazA^min^ CP (see S.I. 2.7 for sequence choices). Expression and modification of these peptides by FIT-Laz and the subsequent LC-MS analysis (Fig. 4a) revealed that 9 out of 10 substrates produced thiopeptides as efficiently as LazA^min^. One substrate (**10aa-sub-4**) led to the formation of multiple thiopeptides owing to three Ser and Thr residues in the inserted region undergoing differential dehydration (**Fig. S16**), and another variant (**10aa-sub-10)** had major accumulation of partially processed linear peptides, albeit with detectable formation of the thiopeptide.

**Figure 4.**
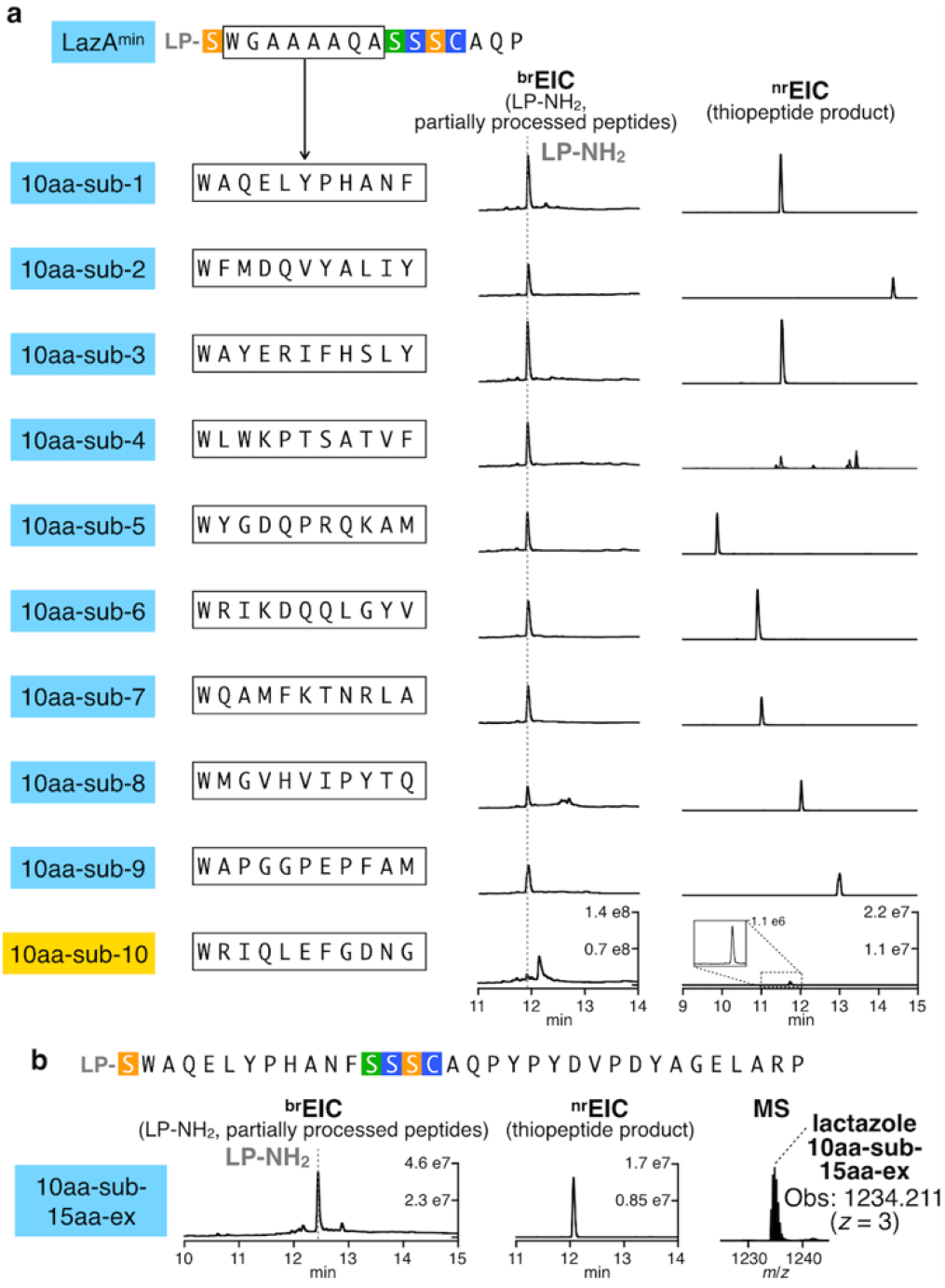
Synthesis of lactazole-like thiopeptides with randomized sequences. **(a)** LazA^min^ variants containing 10 consecutively randomized amino acids were first treated with LazDEF, and then with LazBC, tRNA^Glu^ and GluRS (see S.I. 2.6 for details) and the outcomes were analyzed by LC-MS. Displayed are LC-MS chromatograms (^br^EIC on the left showing partially processed linear peptides and LP-NH_2_ after enzymatic treatment, and ^nr^EIC chromatograms on the right for expected thiopeptides generated at *m/z* 0.10 tolerance window). Mutants highlighted in light blue indicate efficient thiopeptide assembly; in yellow – inefficient thiopeptide formation accompanied by the accumulation of linear intermediates and side-products. One construct, 10aa-sub4, resulted in 8 different thiopeptides, partial annotation of which can be found in **Fig. S16**. Efficient *in vitro* biosynthesis observed in 9 out of 10 cases underscores the substrate plasticity of FIT-Laz. **(b)** LC-MS chromatograms as in (a) for the enzymatic processing of a 34 amino acid-long LazA^min^ variant on the left with a zoomed-in mass spectrum of the produced thiopeptide on the right.

Finally, we combined sequence randomization inside the macrocycle with the C-terminal extension, and constructed a LazA^min^ mutant with a 34 amino acid-long CP. Despite its size, this substrate efficiently generated a 3.7 kDa thiopeptide when treated with Laz enzymes (Fig. 4b, **S17**), highlighting the scaffolding ability of key residues in LazA^min^.

Taken together, these data indicate an unprecedented flexibility of *laz* BGC. Many individual enzymes and entire RiPP pathways are similarly promiscuous, but thiopeptide biosynthesis is usually sensitive to much more modest perturbations. These results point to potential applications of *laz* BGC in bioengineering.

### Synthesis of hybrid thiopeptides with FIT-Laz

One advantage of the FIT system is its amenability to genetic code reprogramming. Incorporation of multiple npAAs can be achieved by adding appropriate orthogonal tRNAs precharged with npAAs of choice by the use of flexizymes^42^ to the translation mixture lacking certain proteinogenic amino acids and cognate aminoacyl-tRNA synthetases. The FIT system was previously used to synthesize peptides containing a variety of npAAs, including D-, β-, *N*-methylated-, and α,α-disubstituted-amino acids as well as hydroxyacids.^62^ Recently, a combination of genetic code reprogramming in the FIT system with a promiscuous RiPP enzyme also enabled synthesis of peptides containing exotic azoline residues.^63^ Such npAAs are often found in peptidic natural products, both in RiPPs^9^ and in non-ribosomally synthesized peptides (**NRP**s).^64^ We reasoned that if LazA precursors containing ribosomally installed npAAs are accepted by Laz enzymes, various “hybrid” thiopeptides may be accessible with the FIT-Laz system.

We began by testing the ability of FIT-Laz to produce *N*-methylated thiopeptides, and prepared 12 *lazA*^*min*^ mutants bearing a single Met codon (AUG) in the CP. The Met codon was reassigned to either *N*-methylglycine (^Me^Gly) or *N*-methylalanine (^Me^Ala) by expressing these genes from a Met-depleted translation mixture in the presence of precharged ^Me^Gly-tRNA_CAU_ or ^Me^Ala-tRNA_CAU_ (see S.I. 2.8 for details). Treatment of these translation products with the full enzyme set (^Me^Gly- and ^Me^Ala-scanning mutagenesis) and the subsequent LC-MS analysis revealed that, similarly to the results of Lys/Glu-scanning, either of the tested *N*-methylated amino acid was easily accepted in 9 positions, while mutations at Trp2, Cys13 and Ala14 were detrimental, affording little to no mature thiopeptide (**Fig. S18**).

Next, we tested whether more diverse npAAs can be incorporated into the thiopeptide scaffold following the same logic. For this study, we focused on *lazA*^*min*^ bearing the AUG codon in position 5, and analogous to the experiments above, prepared LazA^min^ variants containing d-Ala, d-Ser, cycloleucine (cLeu), pentafluorophenylalanine (Phe(F_5_)), 5-hydroxy-tryptophan (Trp(5-OH)), lactic acid (^HO^Ala), β-Met, and β-homoleucine (β-hLeu). All of these substrates were smoothly converted to the corresponding thiopeptides by the action of Laz enzymes, affording hybrid thiopeptides containing a variety of npAAs (Fig. 5a).

**Figure 5.**
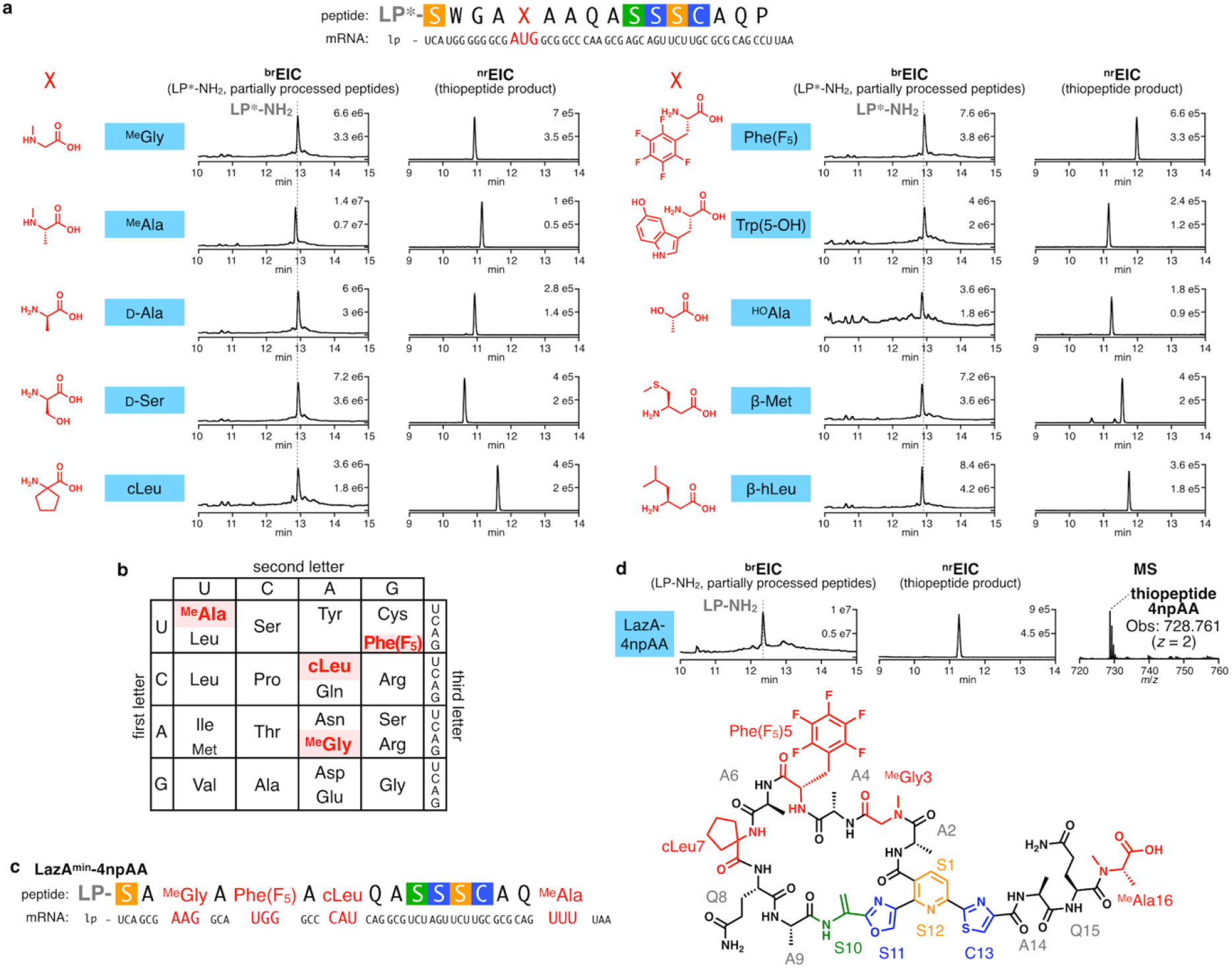
Synthesis of hybrid thiopeptides by genetic code reprogramming with the FIT-Laz system. **(a)** Incorporation of a single npAA in a permissible position 5 of LazA^min^ using the Met AUG codon. LazA^min^ mutants accessed with *in vitro* genetic code reprogramming were treated with the full enzyme set and the reaction outcomes were analyzed by LC-MS. Displayed are LC-MS chromatograms (^br^EIC chromatograms on the left showing partially processed linear peptides and LP*-NH_2_ after enzymatic treatment, and ^nr^EIC chromatograms on the right for expected thiopeptides generated at *m/z* 0.10 tolerance window). LP^*^ stands for LazA LP sequence where formyl-Met is replaced with *N*-biotinylated-Phe (see S.I. 2.8 for details). **(b)** Reprogrammed genetic code utilized for the synthesis of a pseudo-natural lactazole containing 4 npAAs, and **(c)** its mRNA sequence. **(d)** LC-MS chromatograms as in (a) for the enzymatic processing of the LazA^min^ variant from (c), and the chemical structure of the resulting thiopeptide. Taken together, these data suggest that diverse hybrid pseudo-natural thiopeptides are accessible with the FIT-Laz system.

Finally, we studied whether multiple different npAAs can be simultaneously incorporated into the minimal lactazole scaffold to generate highly artificial “pseudo-natural” macrocycles. Due to the presence of a 38-residue LP in LazA^min^, the codon boxes available for reprogramming are limited (see S.I. 2.8 for details). After some experimentation, we opted to reprogram four codons (AAG, CAU, UGG and UUU), and reassigned them as ^Me^Gly, cLeu, Phe(F_5_) and ^Me^Ala, respectively. To this end, a DNA template encoding *lazA*^*min*^ with 4 codons of interest (Fig. 5b and 5c) was incubated in a Lys/His/Phe/Trp-depleted translation reaction with ^Me^Gly-tRNACUU,^Me^Ala-tRNA_AAA_, cLeu-tRNA_GUG_, and Phe(F_5_)-tRNA_CCA_ to yield a LazA precursor peptide bearing 4 npAAs. Treating this substrate with LazBDEF/tRNA^Glu^/GluRS afforded a fully processed linear precursor bearing 2 azoles and 3 Dha (**Fig. S19**), while the reaction utilizing the full enzyme set led to the formation of the predicted thiopeptide accompanied by LP-NH_2_ (Fig. 5d). The identities of this macrocycle and its linear precursor were confirmed by CID MS/MS analysis (**Fig. S19 and S20**). From these experiments, we conclude that the FIT-Laz system offers facile access to previously inaccessible hybrid thiopeptides, including novel heavily modified pseudo-natural architectures. These results additionally underscore the promiscuity of Laz enzymes, as all tested substrates containing disruptive amino acids outside of the canonical Ramachandran space were efficiently converted to mature thiopeptides.

## Discussion

In this study, we completed *in vitro* reconstitution of *laz* BGC, which is responsible for biosynthesis of lactazole A, a cryptic thiopeptide from *S. lactacystinaeus.* This is the first *in vitro* reconstitution of an entire thiopeptide BGC, and the second reconstitution of biosynthetic enzymes involved in the formation of a primary thiopeptide macrocycle.^65^ The FIT-Laz system established in this study enabled rapid access to numerous LazA variants; the entire workflow from PCR assembly of *lazA* DNA templates to LC-MS analysis of reaction outcomes fits within two working days. An added benefit of working with an *in vitro* reconstituted BGC is the ability to decouple self-immunity, export and proteolytic stability issues, so often complicating *in vivo* studies, from direct assaying of enzymatic activities. Conversely, *in vitro* experiments provide no insight into *in vivo* fates and metabolism of the underlying natural product, and thus, should be interpreted accordingly.

The results presented here indicate that *all* Laz enzymes tolerate substantial disruptions in the structure of the precursor peptide (Fig. 6a), which stands in contrast to other thiopeptide BGCs characterized to date. Out of 102 structurally diverse precursor peptides tested in this work, 92 yielded lactazole-like thiopeptides, 73 of which were accessed with efficiencies comparable to wild type lactazole A (Table S8). How the enzymes manage to properly modify such a diverse set of substrates remains to be demonstrated. For now, it is apparent that Laz enzymes are highly cooperative. For instance, addition of LazF to LazDE orchestrates efficient cyclodehydrations (Fig. 2c vs. Fig. 2d), LazC prevents overdehydration by LazBF (Fig. 2h vs. Fig. 2j), and unaminoacylated *S. lactacystinaeus* tRNA^Glu^ somehow affects azole formation mediated by LazDEF in the presence of LazB (**Fig. S6e vs**. Fig. 2d and **S6c–d**). Combined with the fact that some Dha can form independent from azole installation, it is likely that the biosynthetic mechanism is more elaborate than the “azoles form first, Dha second” paradigm observed during thiomuracin biosynthesis^47^ and frequently assumed for other RiPPs. Investigations into the nature of this cooperativity are a subject of our ongoing studies.

**Figure 6.**
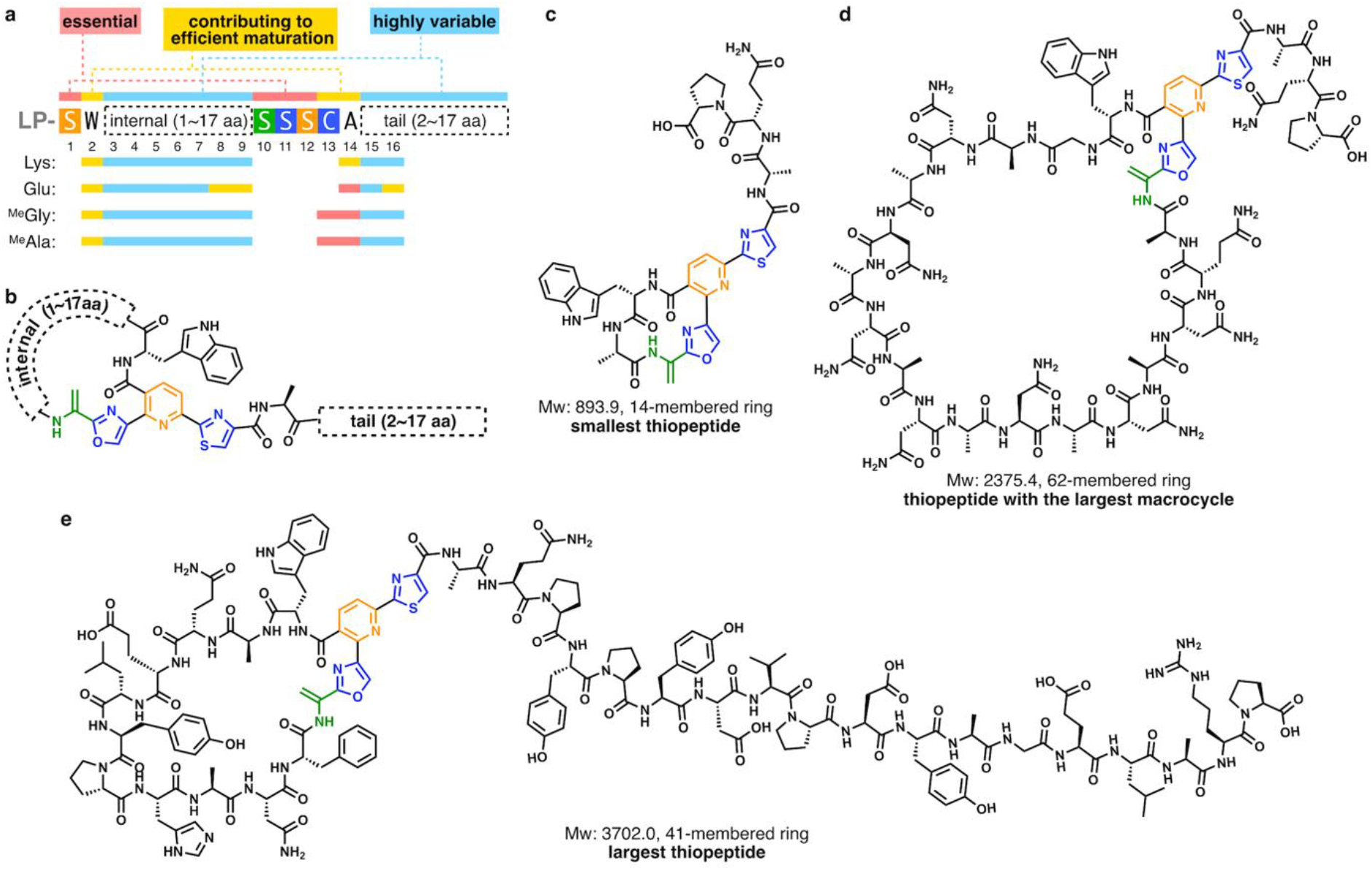
The summary of the work. **(a)** Primary sequence representation of the minimal lactazole scaffold with the outcomes of the Lys, Glu ^Me^Gly and ^Me^Ala scanning experiments mapped to the resulting consensus sequence. **(b)** Chemical structure representation of the minimal lactazole scaffold. **(c)** – **(e)** Structural diversity of thiopeptides accessible with the FIT-Laz system. Displayed are chemical structures of the smallest artificial lactazole (c), thiopeptide with the largest macrocycle (d) and the largest construct (e) synthesized in this work.

Broad substrate scope of Laz enzymes enabled development of the minimal lactazole scaffold (Fig. 6a and b). This thiopeptide requires only 6 PTM events (formation of 2 azoles, 3 Dha and a pyridine heterocycle) for macrocyclization, and is biosynthetically the simplest known thiopeptide to date. The 5 amino acids undergoing these modifications ‒ Ser1, Ser10, Ser11, Ser12 and Cys13 ‒ are indispensable for efficient macrocycle assembly, and further experiments demonstrated that the residues adjacent to the modification sites, Trp2 and Ala14, are also important for efficient biosynthesis. Remaining positions (3–9, 15, 16) accept a variety of amino acids, including disruptive npAAs. Modification of minimal lactazole precursor, LazA^min^, in the FIT-Laz system is robust, and tolerates massive sequence variations. Specifically, the macrocycle can be contracted or expanded to synthesize 14-to 62-membered thiopeptides (2 to 18 unmodified amino acids inside the macrocycle; Fig. 6c and 6d), and the variants with up to 18 amino acid-long tails are accessible as well (Fig. 6e). Most importantly, LazA^min^ can accommodate mutations of consecutive amino acids, as demonstrated by the synthesis of thiopeptides with 10 randomized amino acids inside the macrocycle (Fig. 6e).

This flexibility of the lactazole biosynthetic machinery, combined with the minimal size of *laz* BGC, which contains only the genes essential for macrocyclization, suggest that the minimal thiopeptide scaffold may be an excellent candidate for bioengineering. Because continuous randomized epitopes can be displayed inside a thiopeptide backbone, we envision that combinatorial libraries based on this scaffold can be generated and screened akin to the recent reports on lanthipeptide bioengineering.^12–16^ In those cases, lanthipeptide libraries were prepared with the use of promiscuous lanthipeptide synthases, and could be screened against a protein target of interest with the use of phage/yeast display or with the reverse two-hybrid system. These studies resulted in the discovery of lanthipeptide inhibitors of HIV budding process,^15^ urokinase plasminogen activator^16^ and α_v_β_3_ integrin binders.^13^ Similarly, we anticipate that the integration of the FIT-Laz system with powerful *in vitro* screening techniques such as mRNA display^66^ will provide access to artificial thiopeptides with desirable pharmacological profiles for drug discovery purposes.

Synthesis of thiopeptide hybrids with other RiPP and NRP classes represents another bioengineering avenue explored in this work. Combinatorial biosynthesis is a concept from NRP and PKS fields, where enzymes from different BGCs are combined to act on a single substrate to generate novel natural products.^67,68^ This concept has recently been applied to RiPPs either via simultaneous use of enzymes from near-cognate BGCs^69^ or by devising chimeric LPs,^70^ demonstrating that multiple promiscuous enzymes can act together to produce nonnatural hybrid RiPPs. *In vitro* genetic code reprogramming, easily achievable with FIT-Laz, offers an alternative route to similar hybrids, many of which are inaccessible by existing methods. We demonstrated that thiopeptide-NRP hybrids (macrocycles containing hydroxyacids, d-, β-, *N*-methylated-, and α,α-disubstituted-amino acids), thiopeptide-RiPP hybrids (*N*-methylation and D-amino acids are found in borosins,^71^ lanthipeptides,^54^ proteusins,^72^ phallotoxins^73^ and many other RiPPs families^9^), and thiopeptides with “anthropogenic” amino acids not found in nature (Phe(F_5_) and cLeu) can be routinely accessed with FIT-Laz. Such noncanonical hybrid architectures can further expand the range of available molecular complexity for biotechnology and drug discovery. Overall, we believe that the established FIT-Laz system opens exciting new opportunities for thiopeptide engineering and characterization of natural thiopeptide diversity.

## Acknowledgements

We thank Dr. Kazuya Teramoto, Dr. Takayuki Kuge, Mr. Kakeru Narumi and Dr. Emiko Nagai for their help with protein expression. This work was supported by PRESTO, Japan Science and Technology Agency (JST) to Y.G.; CREST for Molecular Technologies, JST to H.S.; KAKENHI (JP16H06444 to H.S. and Y.G. and H.O.; JP17H04762, JP18H04382, and JP19K22243 to Y.G.) from the Japan Society for the Promotion of Science (JSPS); a grant-in-aid from the Institute for Fermentation, Osaka (IFO) to H.O., S.A. and T.O.; Amano Enzyme, Inc. to H.O and S.A.; and the A3 Foresight Program, JSPS to H.O. and S.A.

