## Supplementary information for "Minimal lactazole scaffold for *in vitro* production of pseudo-natural thiopeptides"

### 1. General

Reagents were purchased from Nacalai Tesque, Wako Pure Chemical Industries, Sigma-Aldrich Japan, Kanto Chemical, or Watanabe Chemical Industries unless noted otherwise and were used as received. Oligonucleotides were purchased from Eurofins Genomics (OPC purification grade) and were used without further purification. *E. coli* DH5α was used for plasmid manipulation. All PCR amplifications were carried out in a BioER TC-96GHBC thermal cycler. Sanger sequencing of constructed plasmids was done with appropriate primers by FASMAC (Tokyo, Japan).

### 2. Methods

#### 2.1. Gene cloning

Primers used in the gene cloning experiments are listed in Table S1, and codon optimized ORFs can be found in S.I. 4. Genes optimized for recombinant expression in *E. coli* were synthesized by Genewiz in pUC57 vectors with *Nde*I and *Xho*I sites flanking each ORF at the 5' and 3' ends, respectively. Cloning procedures followed standard molecular biology techniques.[1] For *lazB* and *lazC*, the codon optimized sequences were cloned into pET26b (Novagen) using *Nde*I/*Xho*I restriction cloning. For *lazD* and *lazE*, the codon optimized sequences were amplified by PCR using appropriate primers, and cloned into pColdII (TAKARA) with *Nde*I/*Bam*HI restriction enzymes. Construction of pET26b-*lazF* was as previously described.[2] The identity of all recombinant constructs was assessed by DNA sequencing, and correct clones were selected for protein expression.

#### 2.2. Protein expression and purification

For LazD and LazE, *E. coli* BL21(DE3) were transformed with the appropriate pCold II cold-shock plasmids using 100 μg/mL carbenicillin as a selection marker, and the resulting transformants were inoculated in Luria-Bertani (LB) medium containing 100 μg/mL carbenicillin. After incubation at 37°C for 16 h, overnight cultures (20 mL) were inoculated in 800 mL LB with 100 μg/mL carbenicillin, and grown at 37°C for 3 hours. The flasks were then transferred on ice, cooled to 4°C, and induced with IPTG to a final concentration of 100 μM. Expression was carried out overnight at 15°C for 20 h while shaking at 180 rpm, and then the cells were harvested by centrifugation (4720 g for 10 min). The pellets were resuspended in 20 mL lysis buffer [50 mM Tris buffer (pH 8.0) containing 500 mM NaCl, 10 mM imidazole and 2 mM DTT], and lysed by sonication on ice using a UD-100 (TOMY) sonicator. The soluble fraction was separated by centrifugation at 4˚C (10300 g for 30 min), filtered, and loaded on a Bio-Scale TM Mini Profinity TM IMAC cartridge preequilibrated with lysis buffer. The column was washed with 50 mL lysis buffer, and then eluted with the same buffer containing 200 mM imidazole (pH 8.0). Fractions containing pure protein were combined, desalted with a Bio-Gel P-6 Desalting Cartridge into protein storage buffer [25 mM HEPES (pH 8.0), 500 mM NaCl, and 2 mM DTT], and concentrated using a 30 kDa cutoff Amicon centrifugal filter (Merck) to 42 μM and 22 μM for LazD and LazE, respectively. Protein concentrations were determined using 280 nm absorbance using extinction coefficients calculated with the ExPASy ProtParam tool (https://web.expasy.org/protparam/). Purity at various purification stages was assessed by Coomassie-stained SDS-PAGE gel analysis. Proteins were flash-frozen and stored at -80°C.

LazB, LazC and LazF were expressed in *E. coli* BL21(DE3) transformed with the appropriate pET26b plasmids. Overnight LB cultures (4 mL) were inoculated in 200 mL of ZYM-5052 autoinducing medium supplemented with 100 μg/mL kanamycin and grown at 18°C for 20 h while shaking at 180 rpm. Purification of these proteins closely followed the protocol described for LazD and LazE, except in these cases lysis buffer lacked DTT, and the final storage buffer was different. For LazB, the storage buffer was 25 mM HEPES (pH 8.0), 500 mM NaCl, 5% glycerol; for LazC and LazF – 25 mM HEPES (pH 6.8), 500 mM NaCl. Final protein stock concentrations were 18 μM for LazB, 23 μM for LazC, and 33 μM for LazF. For LazF, the concentration was determined using a Bradford colorimetric assay.

For *S. lividans* GluRS expression, *E. coli* BL21(DE3) transformed with pET26b-gluRS were inoculated in LB medium containing 100 μg/mL kanamycin and incubated at 37°C for 16 h. Then, 4 mL of the overnight culture was inoculated in 200 mL LB medium containing 100 μg/mL kanamycin as a selection marker, and the cells were grown at 37°C for 2 h while shaking at 150 rpm. The flasks were transferred on ice, and the culture was induced with IPTG to a final concentration of 100 μM. Expression was done at 18˚C for 20 h while shaking at 180 rpm. The purification protocol followed LazD and LazE, except lysis buffer contained 50 mM Tris (pH 8.0), 300 mM NaCl, 20 mM imidazole. After His-tag purification, the protein was dialyzed against 50 mM Tris (pH 7.5). GluRS was concentrated as described above to a final concentration of 36 μM and flash-frozen for storage.

#### 2.3 Preparation of *lazA* synthetic DNA

Linear double-stranded DNA encoding T7 promoter upstream of *lazA* ORF and the mutants of thereof were assembled by PCR from synthetic single-stranded DNA oligonucleotides using *Taq* polymerase. All PCR were performed in 10 mM Tris (pH 8.4), 50 mM KCl, 0.1% (v/v) Triton X-100,

2.5 mM $MgCl_2$, 250 µM each dNTP supplemented with 500 nM of appropriate primers and *Taq* DNA polymerase (standard PCR conditions). Three stage thermal cycling included a denaturing step at 95°C for 40 s, annealing at 52°C for 40 s, and extension at 72°C for 40 s. The list of all oligonucleotides and assembly schemes can be found in Tables S2 and S3.

Briefly, for wild-type *lazA*, forward and reverse primers were annealed, and extended in the primer extension step. To this end, 100 µL PCR solution containing 500 nM primers was denatured at 95°C for 60 s. Then, five cycles of annealing (52°C for 60 s) and extension (72°C for 60 s) were performed, and 1 µL of the product was used as a template for the next step. The first PCR was performed under the standard conditions with 5 cycles of amplification. The second PCR was performed on a 1000 µL scale using 5 µL of the first PCR product with 14 cycles of amplification, and the outcome was evaluated by agarose gel electrophoresis. DNA was first extracted by phenol/chloroform/isoamyl alcohol (25:24:1, saturated with 10 mM Tris (pH 8.0), 1 mM EDTA), and then by chloroform/isoamyl alcohol (24:1). Extracted DNA was precipitated with ethanol, washed with 70% ethanol in water (v/v), and dissolved in 100 µL of water.

Other templates were assembled using wild type or $lazA^{min}$ as a template in one or two steps. For single step mutagenesis, reactions were performed on a 200 µL scale with 1 µL of 1:100 diluted template DNA and 14 cycles of amplification. For two step procedures, 100 µL of PCR solution containing 0.5 µL of 1: 10 diluted template DNA was amplified for 10 cycles, followed by a 14-cycle amplification of the PCR product from the first step (1 µL in 200 µL PCR solution). Analysis of amplification and DNA isolation were performed as above. These DNA templates were used for *in vitro* translation as is, without concentration adjustment.

#### 2.4. Preparation of tRNA and flexizymes

tRNAs and flexizymes were prepared by *in vitro* transcription with T7 RNA polymerase from DNA templates encoding corresponding sequences downstream of a T7 promoter. Analogous to wild type *lazA*, DNA was assembled by PCR. Forward and reverse primers were extended, and then further amplified by PCR under the standard conditions. Primer sequences and assembly schemes can be found in Tables S4 and S5. PCR products were extracted by phenol/chloroform/isoamyl alcohol (25: 24: 1, saturated with 10 mM Tris (pH 8.0), 1 mM EDTA), chloroform/isoamyl alcohol (24: 1), and precipitated with ethanol. DNA were redissolved in water and added to the transcription reaction mix (40 mM Tris buffer (pH 8.0) supplemented with 22.5 mM $MgCl_2$, 10 mM DTT, 1 mM spermidine, 0.01% Triton X-100, 120 nM T7 RNA polymerase, 0.04 U/µL RNasin RNase inhibitor, and 3.75 mM each NTP). For tRNA transcription, 5 mM GMP was additionally supplied to the reaction mixture. Transcriptions were conducted at 37°C for 12-16 h on a 2 mL scale. After, 60 µL of 1 unit/µL RQ1 RNase-free DNase (Promega) was added, and the reactions were further incubated for 60 min at 37°C. The transcripts were precipitated with isopropanol, redissolved in water, and purified by 8% (tRNAs) or 12% (flexizymes) polyacrylamide gel containing 6 M urea. RNA extracted from the gel with 300 mM NaCl were collected by ethanol precipitation followed by centrifugation (15300 g for 15 min), and dissolved in water for storage.

#### 2.5. tRNA Aminoacylation

For genetic code reprogramming experiments, $tRNA^{fMet}$, $tRNA^{GluE2}$ and $tRNA^{AsnE2}$ were aminoacylated with non-proteinogenic amino acids by the use of flexizymes. In general, 25 µM tRNA and 25 µM flexizyme were incubated with 5 mM activated amino acid ester (3,5-dinitrobenzyl or cyanomethyl esters; all prepared as previously established)[3] in 50 mM HEPES-KOH buffer (pH 7.5) containing 600 mM $MgCl_2$ on ice for 2 h. The reactions were stopped with the addition of 300 mM NaOAc (pH 5.2), and precipitated with ethanol. Precipitated RNA was recovered by centrifugation (15300 g for 15 min). The pellets were washed with 70% ethanol in water (v/v) containing 100 mM NaOAc (pH 5.2) and used in translation. For some amino acids, conditions deviated from this general protocol; a complete list of aminoacylation conditions (flexizyme, activated ester, pH, and reaction time) can be found in Table S6.

#### 2.6. In vitro translation and enzymatic reactions

A transcription-coupled *in vitro* translation system was reconstituted as previously reported by mixing purified ribosome, enzymes and translation factors. The final reaction mixture contained 50 mM HEPES-KOH (pH 7.6), 100 mM KOAc, 2 mM GTP, 2 mM ATP, 1 mM CTP, 1 mM UTP, 20 mM creatine phosphate, 12 mM $Mg(OAc)_2$, 2 mM spermidine, 2 mM DTT, 1.5 mg/mL *E. coli* total tRNA (Roche), 1.2 µM ribosome, 0.6 µM MTF, 2.7 µM IF1, 0.4 µM IF2, 1.5 µM IF3, 10 µM EF-Tu, 10 µM EF-Ts, 0.26 µM EF-G, 0.25 µM RF2, 0.17 µM RF3, 0.5 µM RRF, 4 µg/mL creatine kinase, 3 µg/mL myokinase, 0.1 µM pyrophosphatase, 0.1 µM nucleotide-diphosphatase kinase, 0.1 µM T7 RNA polymerase, 0.73 µM AlaRS, 0.03 µM ArgRS, 0.38 µM AsnRS, 0.13 µM AspRS, 0.02 µM CysRS, 0.06 µM GlnRS, 0.23 µM GluRS, 0.09 µM GlyRS, 0.02 µM HisRS, 0.4 µM IleRS, 0.04 µM LeuRS, 0.11 µM LysRS, 0.03 µM MetRS, 0.68 µM PheRS, 0.16 µM ProRS, 0.04 µM SerRS, 0.09 µM ThrRS, 0.03 µM TrpRS, 0.02 µM TyrRS, 0.02 µM ValRS, 500 µM each proteinogenic amino acid and 100 µM 10-HCO-H4 folate. Translations were performed at 37°C for 50 to 60 min with 1 μL of *lazA* variant template DNA, for a total translation volume of 5 μL.

The translation product was split in two equal parts. One part was treated with 12.5 μL of the Laz enzyme mix, which contained 2 μM LazB, 2 μM LazC, 1 μM LazD, 1 μM LazE, 2 μM LazF, 1 μM *S. lividans* GluRS, 10 μM *S. lactacystinaeus* $tRNA^{Glu}$ in 50 mM Tris buffer (pH 8.0) supplemented with 10 mM $MgCl_2$, 5 mM ATP, and 1 mM DTT. The second half was incubated with the same buffer lacking enzymes and tRNA, as a translation control. After a 15-16 h incubation at 25°C, the reactions were transferred on ice, and 1 volume of 30 mM iodoacetamide in methanol was added. Precipitated protein and nucleic acid were separated by centrifugation (15300 g for 4 min), and 10 μL of the supernatant was analyzed by LC-MS.

Randomized peptides were modified via a two-step protocol. Translation product (2.5 μL) was first incubated with LazDEF for 6 h, and then LazBC, $tRNA^{Glu}$ and GluRS were added to final concentrations as specified above. After a 12 h incubation, sample preparation and analysis followed the general protocol.

#### 2.7. Design of randomized sequences

Randomized peptide sequences were generated with ExPASy RandSeq tool (https://web.expasy.org/randseq/), using average amino acid compositions computed from Swiss-Prot. Peptides containing Cys, overly repetitive or hydrophobic sequences were discarded. Negatively charged amino acids in positions 3, 4, 11 and 12 (sites adjacent to the residues undergoing PTM), and positively charged amino acids in positions 11 and 12 were avoided as well.

#### 2.8 *In vitro* translation with genetic code reprogramming.

Due to the presence of a 38-residue LP in LazA composed of 16 kinds of proteinogenic amino acids, the codon boxes available for reprogramming are limited to His, Lys, Phe and Tyr codons. Additionally, the AUG Met codon can be utilized if translation initiation is also reprogrammed, and the UGG Trp codon is available if a mutation of Trp2 in the CP is deemed acceptable. We chose the AUG codon for incorporation of single npAAs, and UGG, CAU, AAG and UUU codons for synthesizing a thiopeptide with 4 npAAs.

Genetic code reprogramming experiments generally followed the workflow from above, except aminoacylated tRNA replaced specific proteinogenic amino acids corresponding to the reprogrammed codons during translation. For incorporation of single npAAs, 50 μM *N*-biotinylated-Phe-$tRNA^{fMet}_{CAU}$ and 50 μM npAA-$tRNA^{GluE2}_{CAU}$ were added to the translation mixture that lacked Met and 10-HCO-H4 folate. As a result, peptides bearing *N*-biotinylated-Phe instead of formyl-Met, and an npAA instead of Met were expressed. Analogously, expression from a Lys/His/Phe/Trp-depleted translation reaction supplemented with $^{Me}$Gly-tRNA$^{GluE2}_{CUU}$, $^{Me}$Ala-tRNA$^{GluE2}_{AAA}$, cLeu-tRNA$^{GluE2}_{GUG}$, and Phe(F$_5$)-tRNA$^{AsnE2}_{CCA}$ resulted in the peptide containing 4 npAAs of interest. In this case, the concentrations of exogenously added tRNA were adjusted to 25 μM each.

#### 2.9. LC-MS analysis of enzymatic reactions

Waters Xevo G2-XS QTof instrument equipped with Acquity I-Class UPLC system was used for LC-MS analysis. HPLC was done on an Acquity UPLC Peptide BEH C18 column (dimensions: 150 x 2.1 mm; pore size: 300 Å; particle size: 1.7 μm) using 0.1% (v/v) formic acid in water (solvent A) or acetonitrile (solvent B) as a mobile phase. Analysis was performed at 60°C and 250 μL/min flow rate running the following gradient: 1% B for 2 min; 1 to 81% B over 20 min; 95% B for 2 min; 1% B for 6 min (total run time: 30 min). MS analysis was done in a positive polarity/high sensitivity mode with a 0.3 s scan time. Capillary voltage was set to 700 V; ESI source and desolvation temperatures were 120 and 400°C, respectively. Manufacturer-supplied GFB was used as a lockspray standard for continuous mass axis referencing, and the recommended lockspray setup procedure was performed prior to every run.

For tandem mass spectrometry, a data-dependent acquisition method was used. CID fragmentation was triggered in real time if detected ions met the following conditions: total ion intensity exceeds $4{\cdot}10^4$ ions, and the ion charge was equal to 3, 4 or 5 (for analyses involving linear LazA precursors) or $z$=2 (for thiopeptides). MS/MS spectra were acquired with a 2 s scan time, with parameterized collision energy values. In method 1, collision energies were set to ramp from 6-8 to 30-40 eV over the range of acquired $m/z$ values (200 to 2000), and in method 2, these values were 6-8 to 45-55 eV. Both methods were utilized for each analyzed peptide, and a more informative spectrum was carried forward with. LC-MS data was analyzed with MassLynx v.4.1, and MS/MS assignments were done in Unifi v.1.8.2.169, allowing relevant PTMs where needed. Fragmentation assignments for thiopeptides were done manually.

#### 2.10. LC-MS data analysis

Broad extracted ion current (broad EIC, $^{br}$EIC) chromatograms were found to be the most informative way to visualize reaction progress (**Fig. S2–S5**). Despite methanol precipitation of translation proteins, nucleic acids and Laz enzymes, significant interference in total ion current (TIC) chromatograms was observed, but in general, most interfering compounds had low molecular weight (<2000 Da), and thus, the $m/z$ region above 1000 stayed relatively clean over the course of a run.

Linear LazA precursors, modified or not, were predominantly detected as $z$=4 species, and as such, had $m/z$ values well above 1000. Generating EIC chromatograms for LazA analogs at $z$=4 with $m/z$ ±100 (±400 Da) tolerance window enabled visualization of reaction outcomes without interference from the translation components. With the exception of the final macrocyclization reaction, all linear intermediates and side-products had mass changes less than the indicated ±400 Da. LP-$NH_2$ also fell within the resulting EIC range at $z$=3, and thus, $^{br}$EIC chromatograms captured all detectable modifications on LazA, and were used to judge reaction outcomes in a semi-quantitative fashion. For each reaction, the validity of this procedure was cross-checked against manual inspection of a parent TIC chromatogram.

The second product of the macrocyclization reaction, a thiopeptide, was outside of this range and had to be visualized separately. For thiopeptides, narrow range EIC ($^{nr}$EIC) chromatograms were generated with an $m/z$ tolerance window generally set to 0.10. When comparing ion intensities of thiopeptides against each other, charge state series EIC traces were summed to yield a full intensity chromatogram for each relevant thiopeptide.

A summary of observed and calculated molecular weights for all discussed compounds can be found in Table S7.

#### 3. Supporting Figures

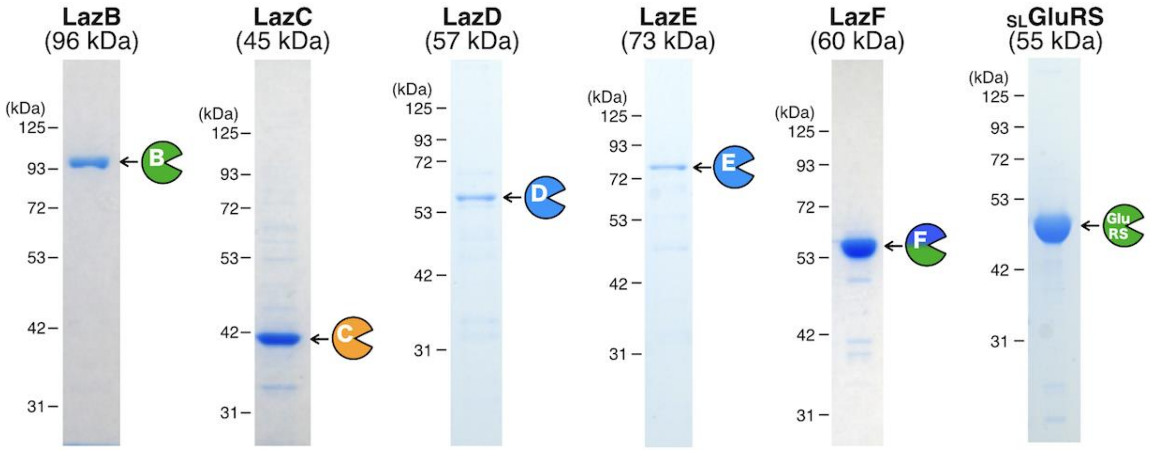

**Figure S1.** SDS-PAGE analysis of recombinantly produced lactazole biosynthetic enzymes utilized in this work. For protein expression details refer to S.I. 2.2.

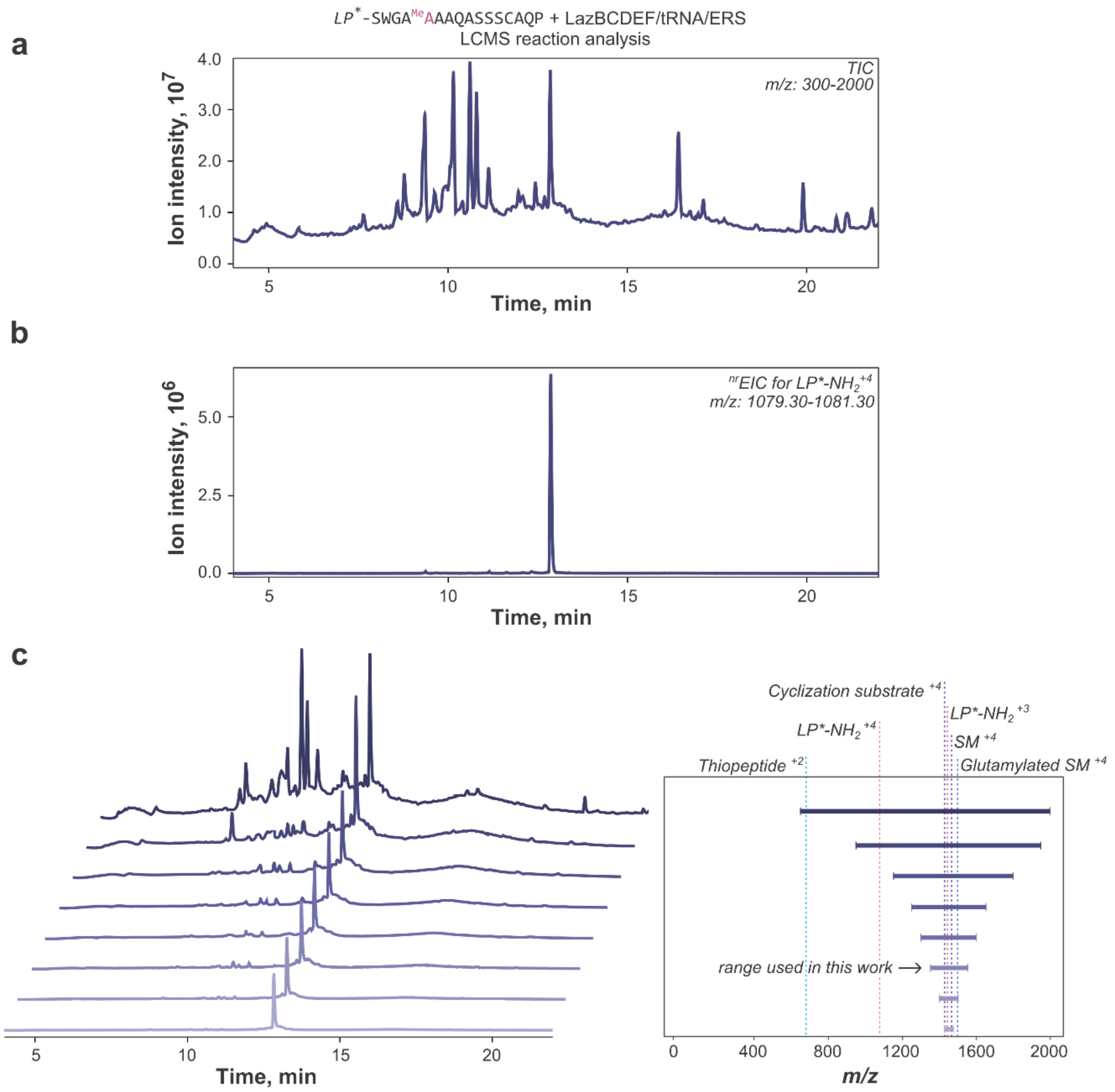

**Figure S2**. An illustration to the data analysis routines and the $^{br}$EIC concept described in S.I. 2.10. **(a)** TIC chromatogram for the reaction between a LazA$^{min}$ variant (CP sequence: SWGA$^{Me}$AAAQASSSCAQP) and Laz enzymes. A number of translation components obfuscate the chromatogram. **(b)** EIC at *m/z* 1080.30±1.00 corresponding to LP$^*$-$NH_2$ produced during the final macrocyclization reaction. LP* stands for LazA LP sequence where formyl-Met is replaced with N-biotinylated-Phe (see S.I. 2.8 for details). **(c)** A visual explanation of the $^{br}$EIC concept. With the exception of the final thiopeptide, all linear intermediates and side-products cluster around m/z ~1400, whereas translation components are characterized by lower *m/z* values. Sequential generation of EIC chromatograms with narrower and narrower *m/z* windows eliminates more and more translation components unrelated to the reaction in question. EIC chromatograms at *m/z*(precursor peptide) ± 100 capture all LazA$^{min}$-derived products with the exception of the mature thiopeptide, for which a separate, narrow range chromatogram was generated in each case.

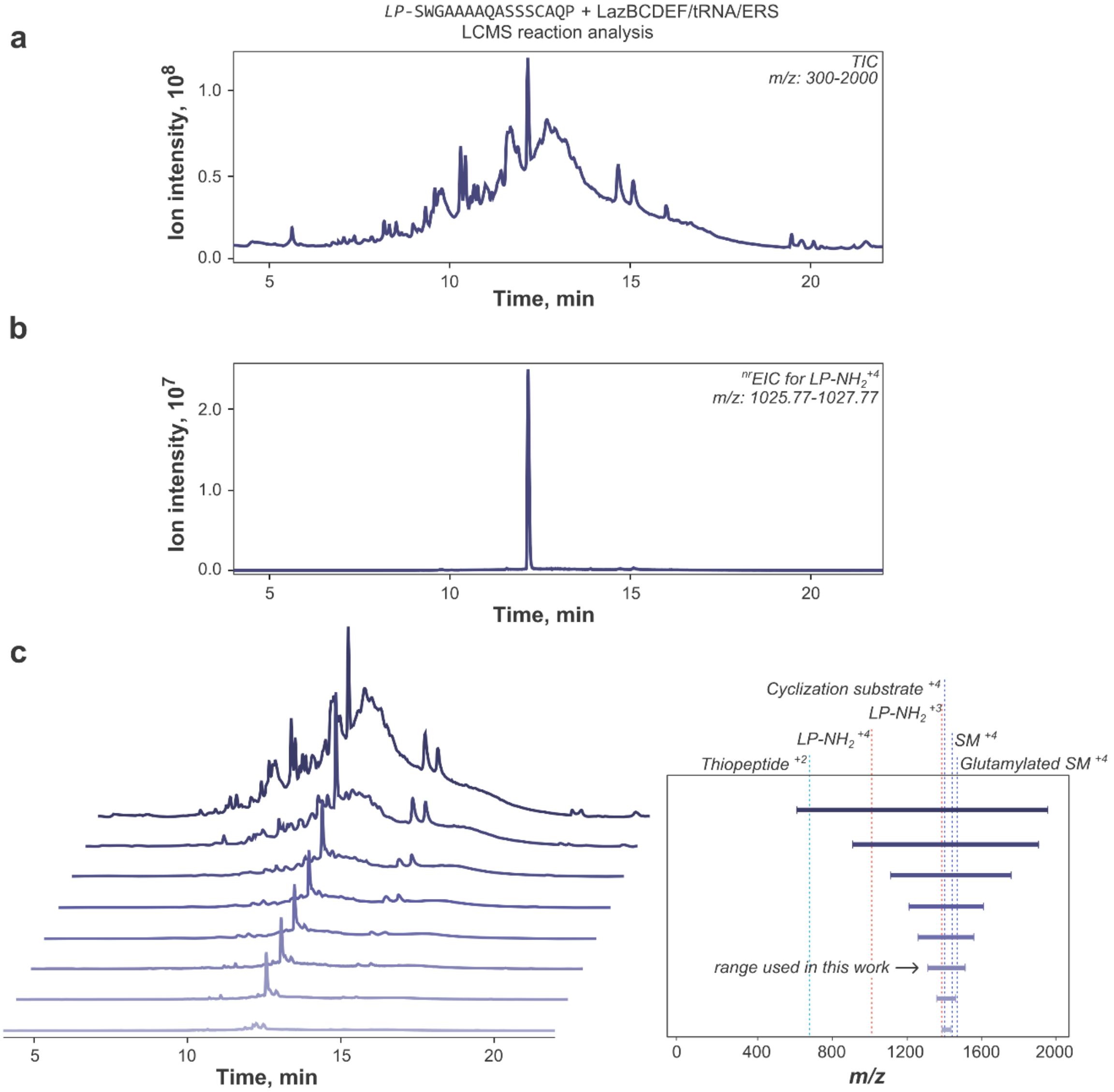

**Figure S3**. An illustration to the data analysis routines and the $^{br}$EIC concept described in S.I. 2.10. **(a)** TIC chromatogram for the reaction between LazA$^{min}$ and Laz enzymes. A number of translation components obfuscate the chromatogram. **(b)** EIC at *m/z* 1026.77±1.00 corresponding to LP-$NH_2$ produced during the final macrocyclization reaction. **(c)** A visual explanation of the $^{br}$EIC concept. With the exception of the final thiopeptide, all linear intermediates and side-products cluster around m/z ~1400, whereas translation components are characterized by lower *m/z* values. Sequential generation of EIC chromatograms with narrower and narrower *m/z* windows eliminates more and more translation components unrelated to the reaction in question. EIC chromatograms at *m/z*(precursor peptide) ± 100 capture all LazA$^{min}$-derived products with the exception of the mature thiopeptide, for which a separate, narrow range chromatogram was generated in each case.

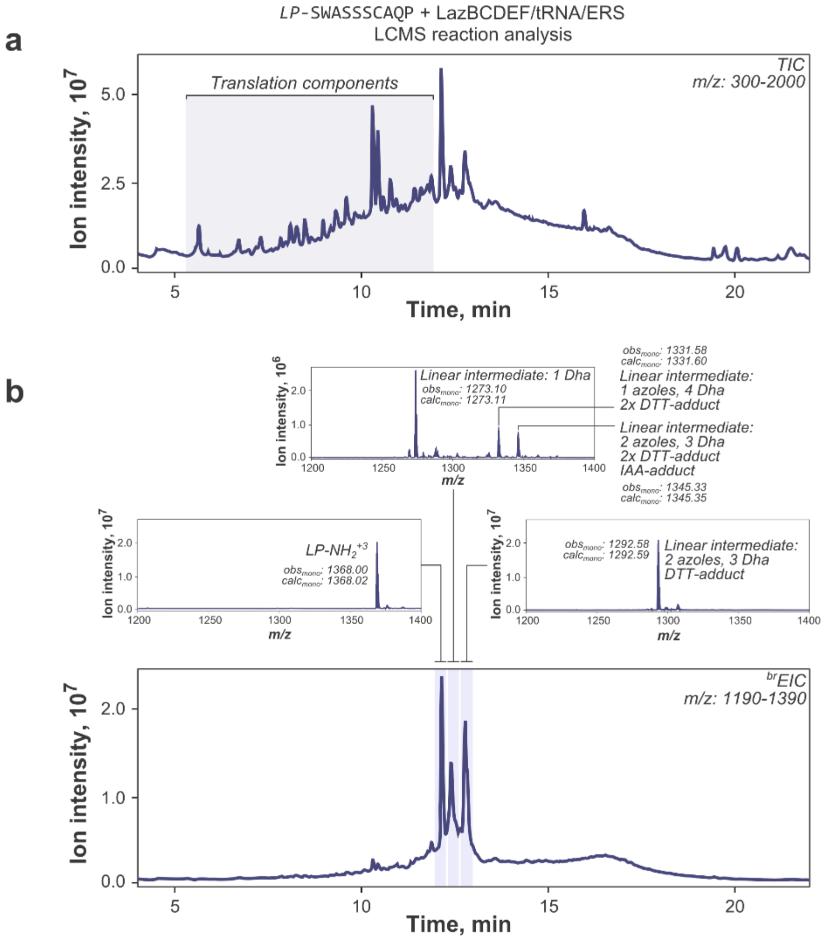

**Figure S4**. An example of LC-MS data analysis performed with the use of brEIC. **(a)** TIC chromatogram for the reaction between a LazA$^{min}$ variant (CP sequence: SWASSSCAQP) and Laz enzymes. A number of translation components obfuscate the chromatogram. **(b)** brEIC chromatogram generated at *m/z* 1290 ± 100 with MS insets integrated over shaded regions. The resulting brEIC chromatogram isolates LazA-related products from translation components, greatly simplifying reaction interpretation and analysis. In this case, the product relevant to thiopeptide formation (LP-$NH_2$; the leftmost peak) is accompanied by a number of linear intermediates and side-products.

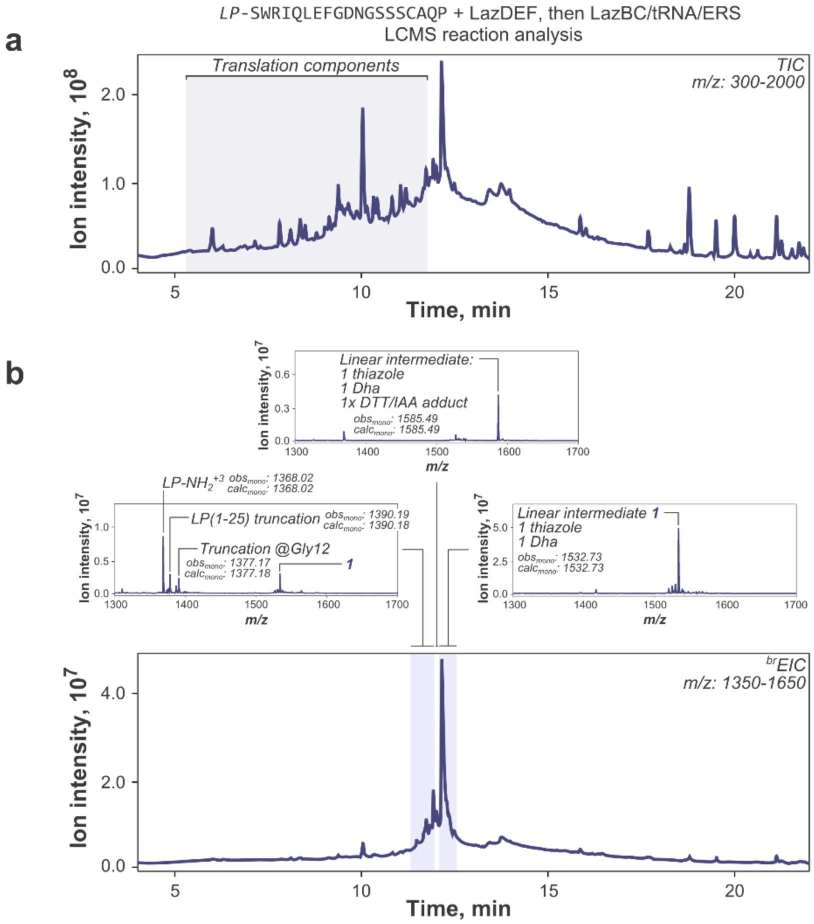

**Figure S5**. An example of LC-MS data analysis performed with the use of $^{br}$EIC. **(a)** TIC chromatogram for the reaction between a LazA$^{min}$ variant (CP sequence: SWRIQLEFGDNGSSSCAQP) and Laz enzymes. A number of translation components obfuscate the chromatogram. **(b)** $^{br}$EIC chromatogram generated at *m/z* 1500 ± 150 with MS insets integrated over shaded regions. The resulting chromatogram isolates LazA-related products from translation components, greatly simplifying reaction interpretation and analysis. In this case, a minute amount of LP-$NH_2$ is accompanied by a number of linear intermediates, side-products and sequence truncations.

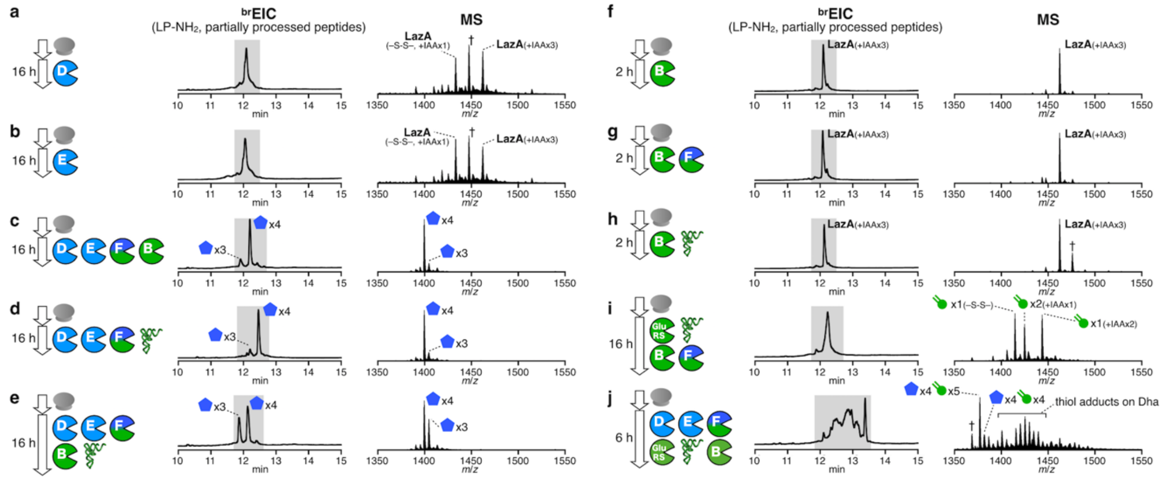

**Figure S6**. Reconstitution of *in vitro* lactazole A biosynthesis. **(a) – (j)** Reconstitution of azole and Dha formation in FIT-Laz. LazA precursor peptide produced with the FIT system was treated with a combination of Laz enzymes as indicated in each panel and the reaction outcomes were analyzed by LC-MS. Displayed are $^{br}$EIC LC-MS chromatograms and composite mass spectra integrated over a time period shaded in the corresponding chromatograms. See S.I. 2.6, S.I. 2.10 and Fig. S2-S5 for details on reaction conditions and the explanation of $^{br}$EIC chromatograms. MS peaks labeled with † are unidentified.

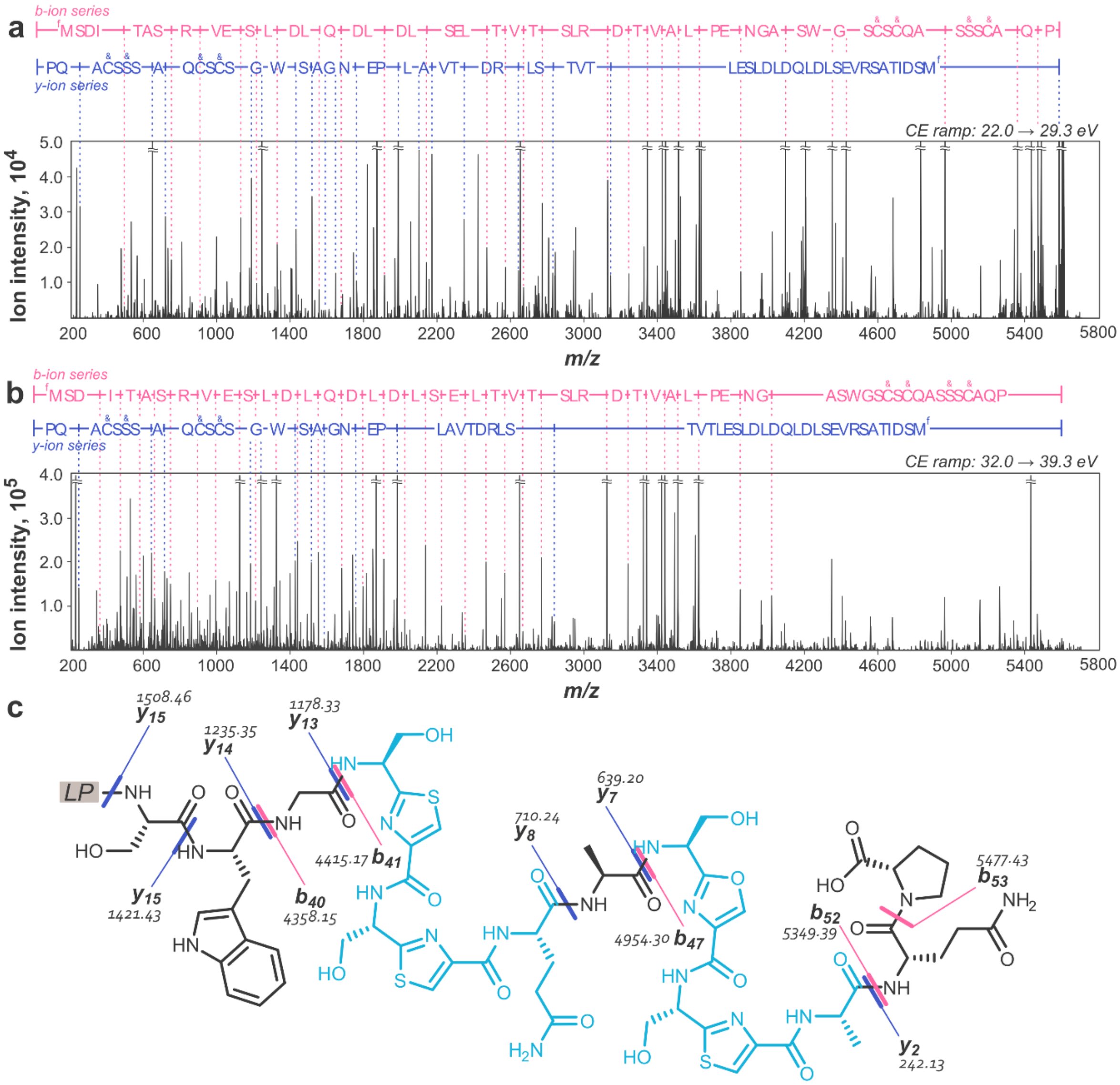

**Figure S7**. MS/MS spectrum of LazA after treatment with LazDEF. **(a)** Charge-deconvoluted CID fragmentation spectrum obtained with collision energies ramped from 22.0 to 29.3 eV and spectral assignments; *y*- and *b*-ions are annotated; stable molecule losses ($H_2O$, $NH_3$, CO, etc) and double fragmentation assignments are omitted for clarity; ampersands denote azole formations; “fM” stands for formyl-methionine; **(b)** Analogous to (a) except the spectrum was obtained with collision energies ramped from 32.0 to 39.3 eV. c) Assigned chemical structure of the LazA CP region with mapped annotations. Each of the two regions highlighted in cyan contains 2 azole modifications, but the precise azole location could not be unambiguously assigned based on these spectra.

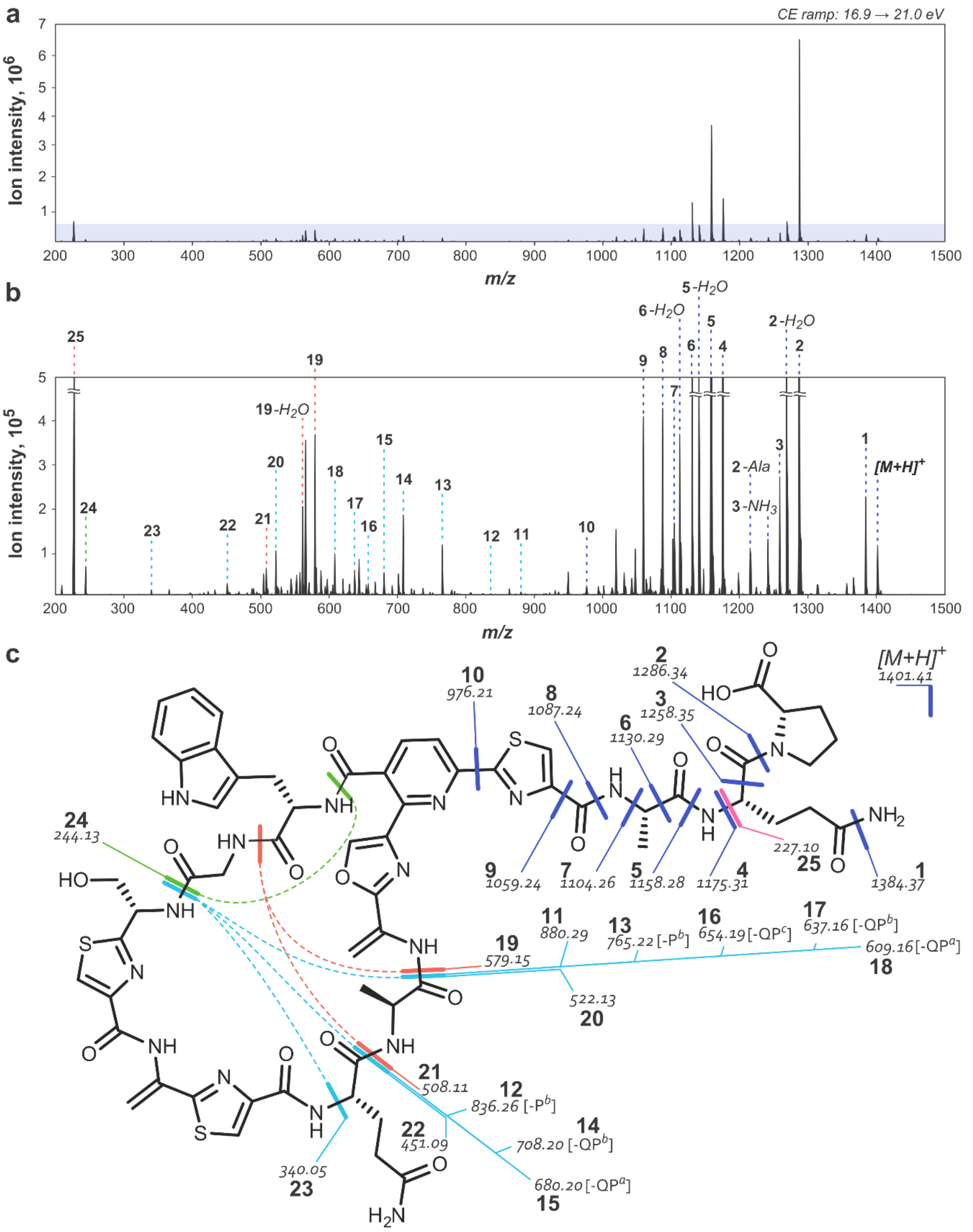

**Figure S8**. Annotated fragmentation spectrum of lactazole A. **(a)** Charge-deconvoluted CID fragmentation spectrum obtained with collision energies ramped from 16.9 to 21.0 eV. **(b)** Y-axis zoom of the shaded area from (a) with spectral assignments. **(c)** Assigned chemical structure of lactazole A with mapped assignments. Under acquisition conditions the thiopeptide underwent multiple double fragmentations allowing the mapping of amino acids within the macrocycle. A number of triple fragmentations are annotated in cyan; for such assignments, positions of the third fragmentation in the tail region are indicated next to the corresponding *m/z* values. Some ions can have multiple potential assignments. In such cases, only 1 isomer is shown.

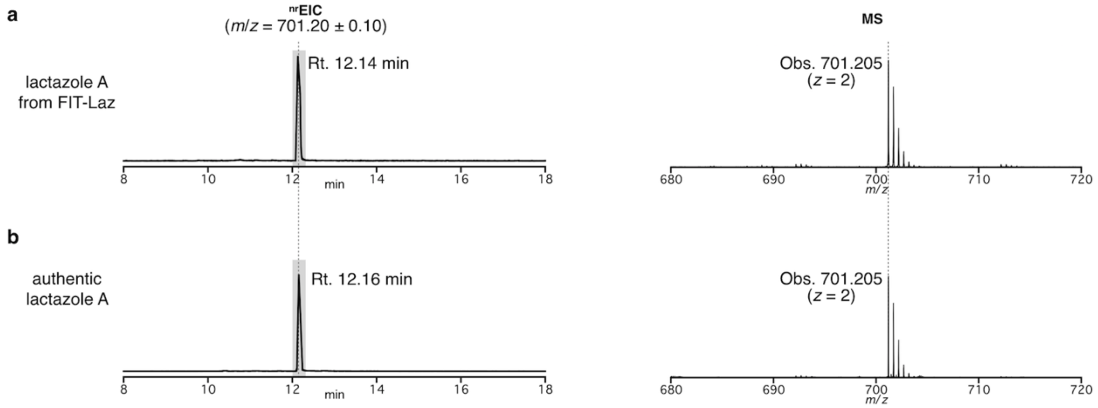

**Figure S9**. LC-MS comparison between authentic lactazole A and lactazole A synthesized with the FIT-Laz system. Left: $^{nr}$EIC chromatograms generated at m/z 701.20 ± 0.10; right: composite mass spectra integrated over a time period shaded in the corresponding chromatograms.

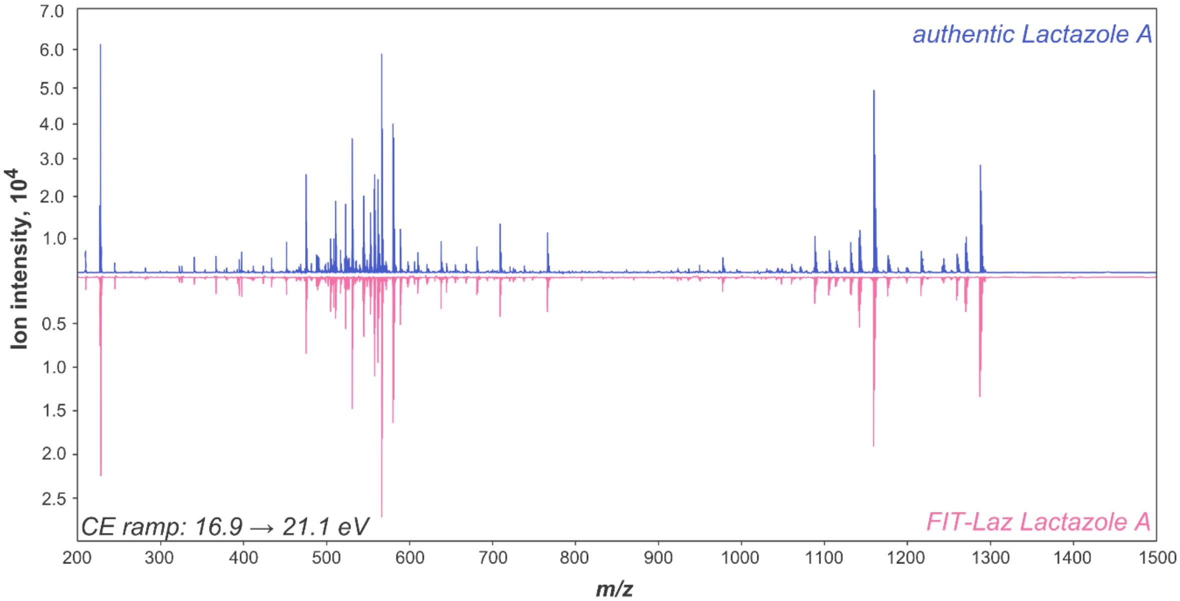

**Figure S10.** Overlaid CID fragmentation spectra of authentic lactazole A (blue spectrum) and lactazole A synthesized with the FIT-Laz system (red). Identical fragmentation patterns confirm authenticity of the synthesized thiopeptide.

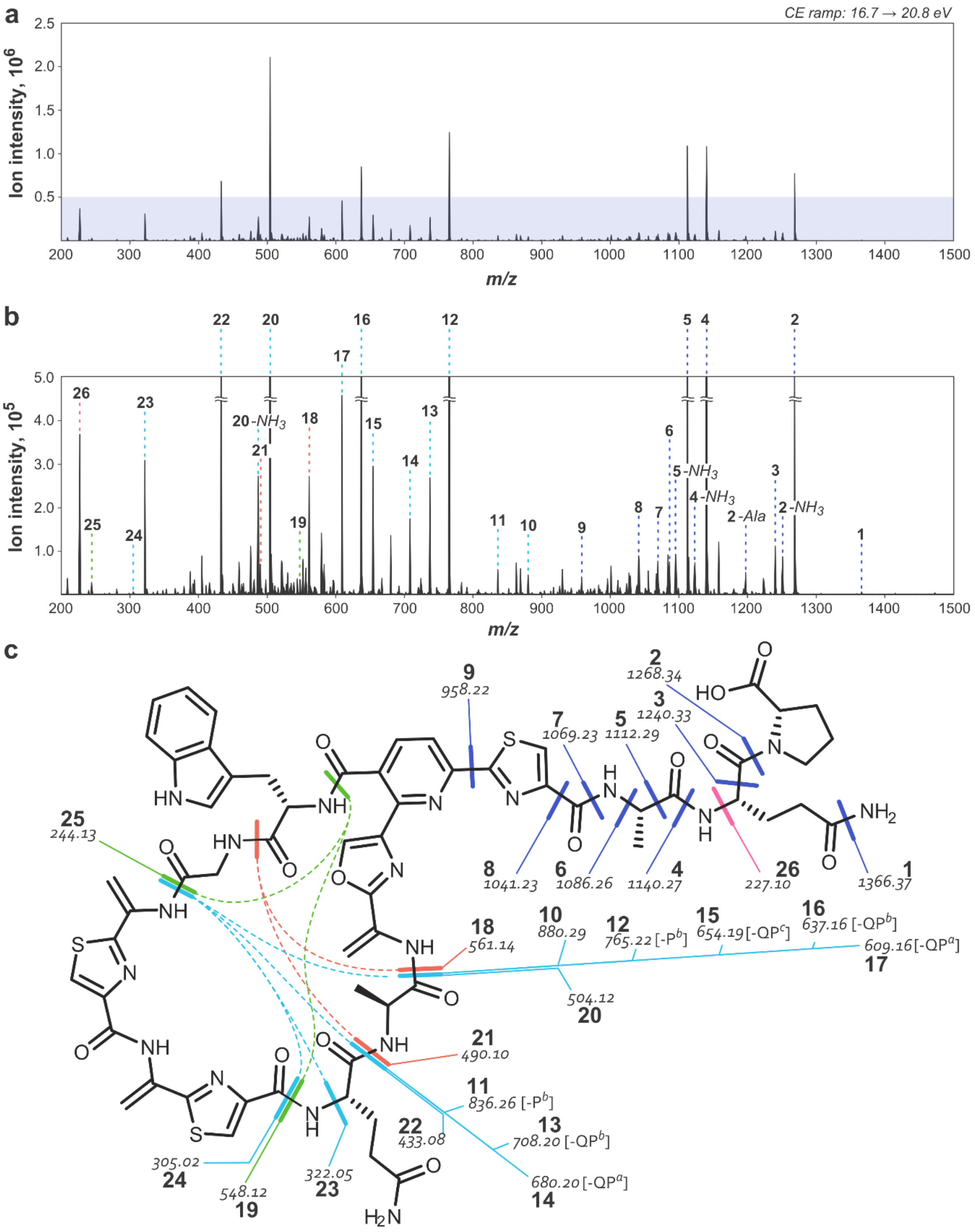

**Figure S11**. Annotated fragmentation spectrum of Dha4-lactazole synthesized with the stepwise enzyme treatment. **(a)** Charge-deconvoluted CID fragmentation spectrum obtained with collision energies ramped from 16.7 to 20.8 eV. **(b)** Y-axis zoom of the shaded area from (a) with spectral assignments. **(c)** Assigned chemical structure of Dha4-lactazole with mapped assignments. Under acquisition conditions the thiopeptide underwent multiple double fragmentations allowing the mapping of amino acids within the macrocycle. A number of triple fragmentations are annotated in cyan; for such assignments, positions of the third fragmentation in the tail region are indicated next to the corresponding *m/z* values. Some ions can have multiple potential assignments. In such cases, only 1 isomer is shown. Extra dehydration compared to lactazole A can be unambiguously localized between residues 4 and 7. Because numerous fragments originating at Gly3-Ser4 bond were observed, dehydration to Dha rather than oxazoline formation at Ser4 can be deduced.

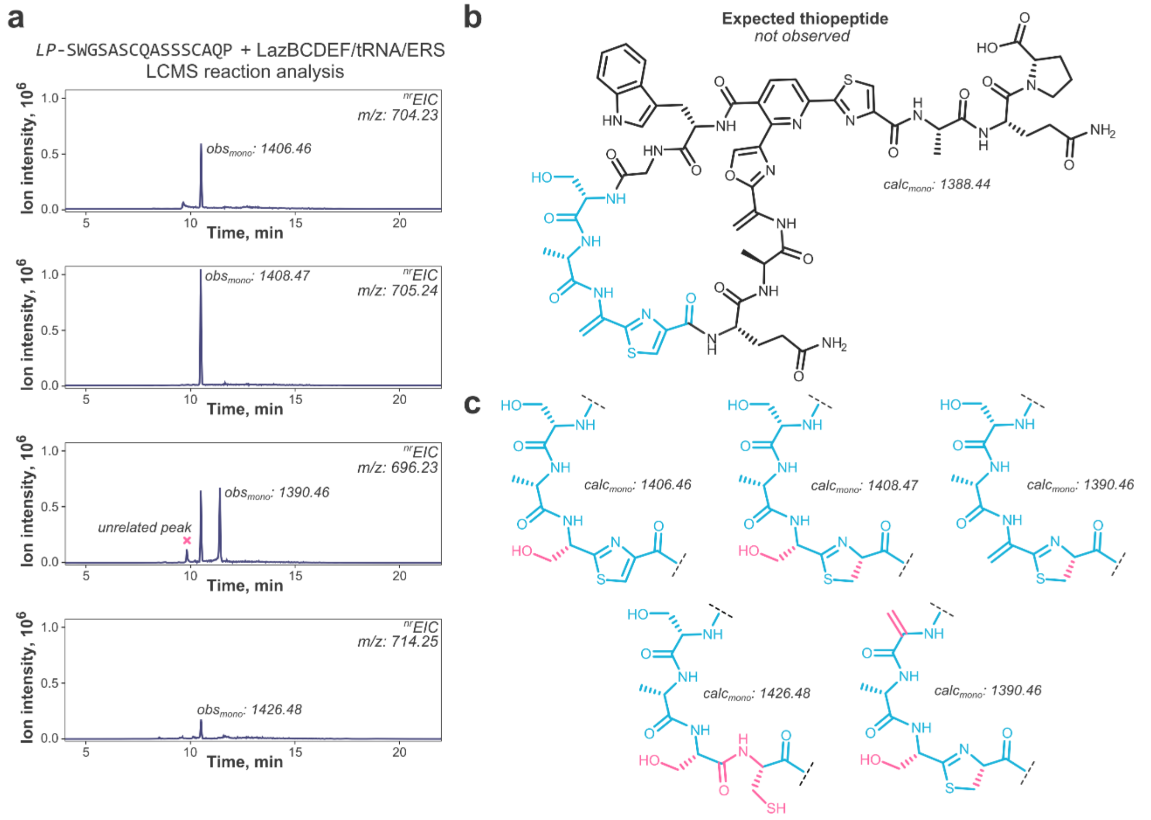

**Figure S12**. Different thiopeptides generated in the FIT-Laz system for LazA C5A mutant. **(a)** $^{nr}$EIC chromatograms generated at *m/z* 0.10 tolerance windows for detected thiopeptides; a total of 4 thiopeptides was observed. **(b)** Chemical structure of the expected lactazole mutant. In this case, the expected product was not detected. The region highlighted in cyan indicates a region with a divergent pattern of PTMs. **(c)** Plausible chemical structures of observed thiopeptides. The constant region is omitted for clarity. Differences between the expected and assigned structures are highlighted in pink.

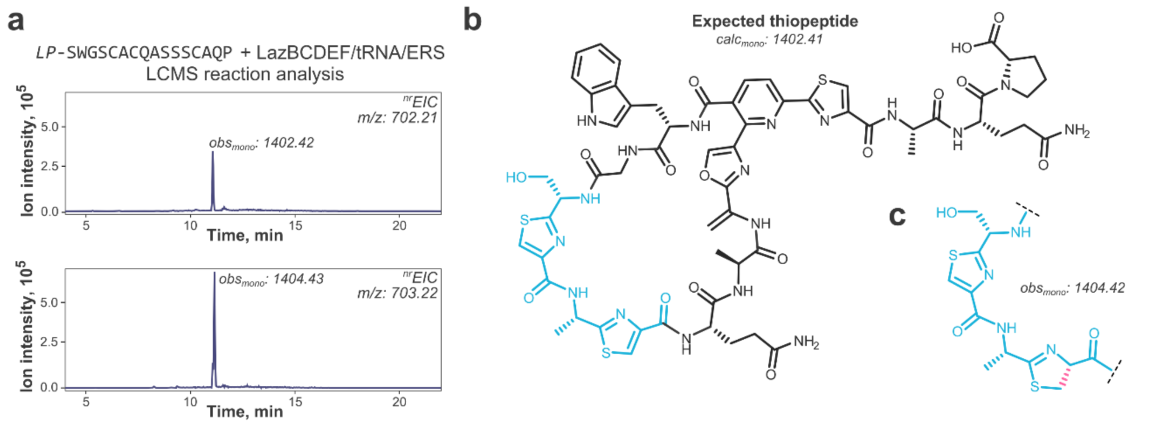

**Figure S13**. Different thiopeptides generated in the FIT-Laz system for LazA S6A mutant. **(a)** $^{nr}$EIC chromatograms generated at *m/z* 0.10 tolerance windows for detected thiopeptides; 2 thiopeptides were observed. **(b)** Chemical structure of the expected lactazole mutant. The region highlighted in cyan indicates a region with a divergent pattern of PTMs. **(c)** Plausible chemical structure observed for the second thiopeptide. The constant region is omitted for clarity. Differences between the expected and assigned structures are highlighted in pink.

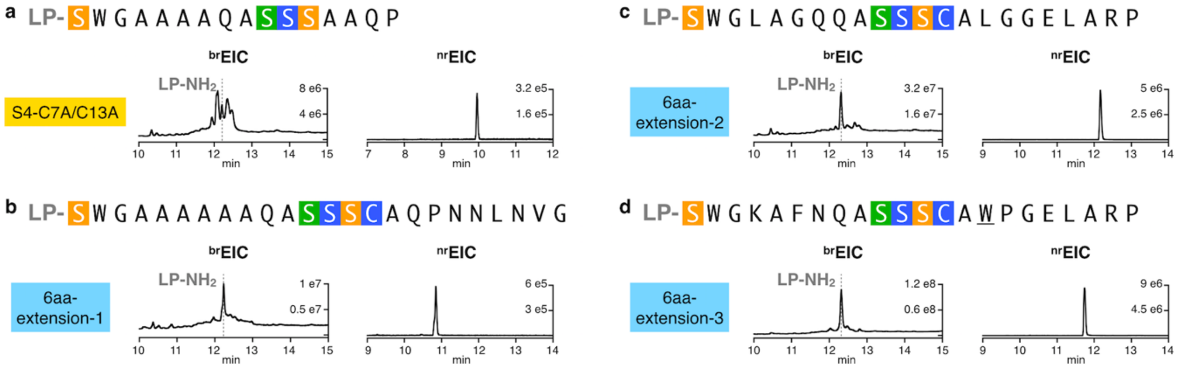

**Figure S14**. Substrate scope of the FIT-Laz system. **(a)** – **(d)** Variants of LazA$^{min}$ were treated with the full enzyme set and the outcomes were analyzed by LC-MS. Displayed are LC-MS chromatograms ($^{br}$EIC chromatograms on the left showing partially processed linear peptides and LP-$NH_2$ after enzymatic treatment, and $^{nr}$EIC chromatograms on the right for expected thiopeptides generated at *m/z* 0.10 tolerance window). For mutants highlighted in light blue biosynthesis proceeded efficiently; yellow highlighting indicates inefficient thiopeptide formation accompanied by the accumulation of linear intermediates and side-products.

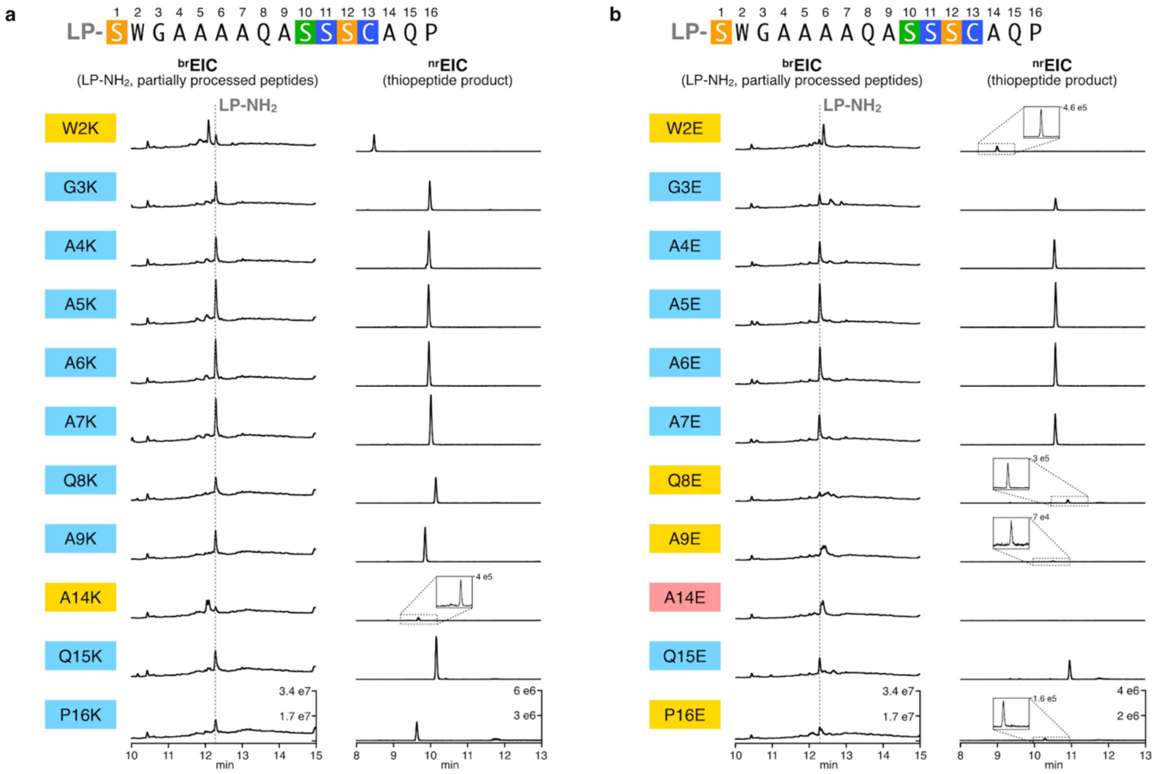

**Figure S15**. Tolerance of Laz enzymes towards charged amino acids in the CP of LazA$^{min}$. **(a)** Lys-scanning and **(b)** Glu-scanning mutagenesis of LazA$^{min}$. Precursor peptides were treated with the full enzyme set and the outcomes were analyzed by LC-MS. Displayed are LC-MS chromatograms ($^{br}$EIC chromatograms on the left showing partially processed linear peptides and LP-$NH_2$ after enzymatic treatment, and $^{nr}$EIC chromatograms on the right for expected thiopeptides generated at *m/z* 0.10 tolerance window). For mutants highlighted in light blue biosynthesis proceeded efficiently; yellow highlighting indicates inefficient thiopeptide formation accompanied by the accumulation of linear intermediates and side-products; red – mutants that failed to yield a detectable thiopeptide.

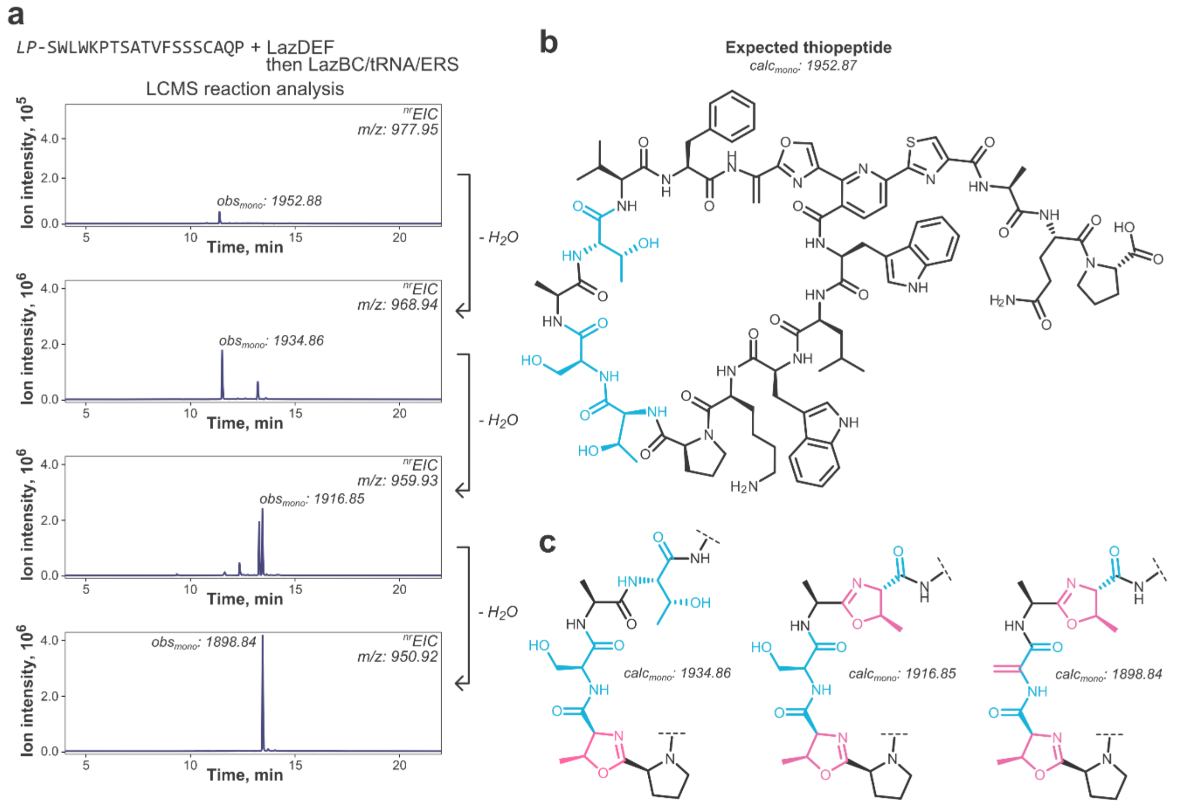

**Figure S16**. Different thiopeptides generated in the FIT-Laz system for LazA$^{min}$ 10aa-sub4. **(a)** $^{nr}$EIC chromatograms generated at *m/z* 0.10 tolerance windows for detected thiopeptide; a total of 8 thiopeptides, differing on their dehydration patterns was observed. **(b)** Chemical structure of the expected thiopeptide. The region highlighted in cyan indicates residues may undergo dehydration in the FIT-Laz system. **(c)** Three plausible chemical structures for sequentially dehydrated thiopeptides. The constant region is omitted for clarity. Differences between the expected and assigned structures are highlighted in pink. Other isomers may arise due to the positional and/or Dha/oxazoline isomerism.

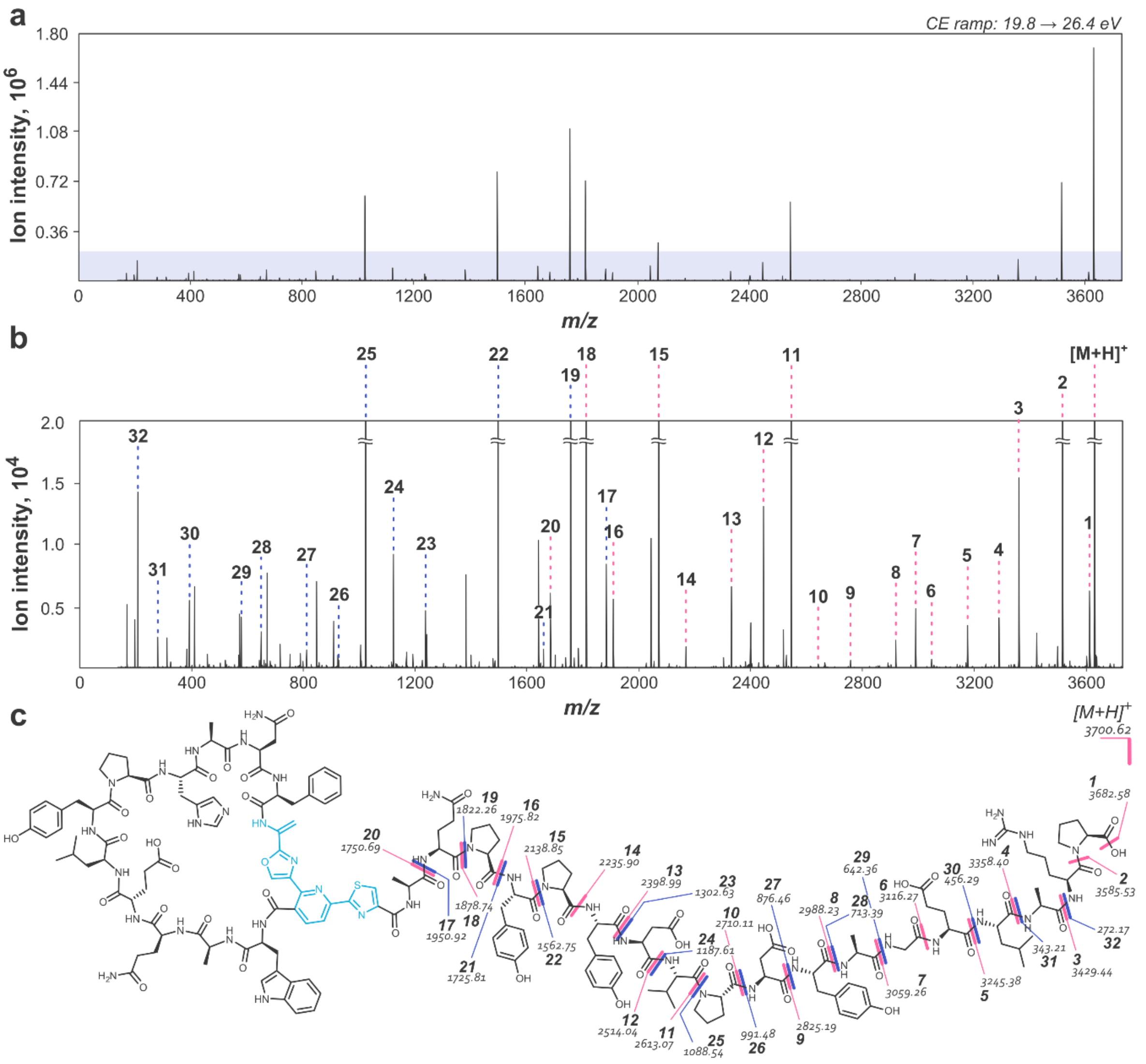

**Figure S17**. Annotated fragmentation spectrum of a 34 amino acid-long pseudo-lactazole synthesized with the FIT-Laz. **(a)** Charge-deconvoluted CID fragmentation spectrum obtained with collision energies ramped from 19.8 to 26.4 eV. **(b)** Y-axis zoom of the shaded area from (a) with spectral assignments; *y*- and *b*-ions are annotated; stable molecule losses ($H_2O$, $NH_3$, CO, etc) and double fragmentation assignments are omitted for clarity. **(c)** Assigned chemical structure of the hybrid thiopeptide with mapped assignments. Under these fragmentation conditions, an almost complete *b*/*y*-fragmentation ladder in the tail region was observed, but no double fragmentation (needed to confirm amino acid sequence inside the macrocycle) occurred.

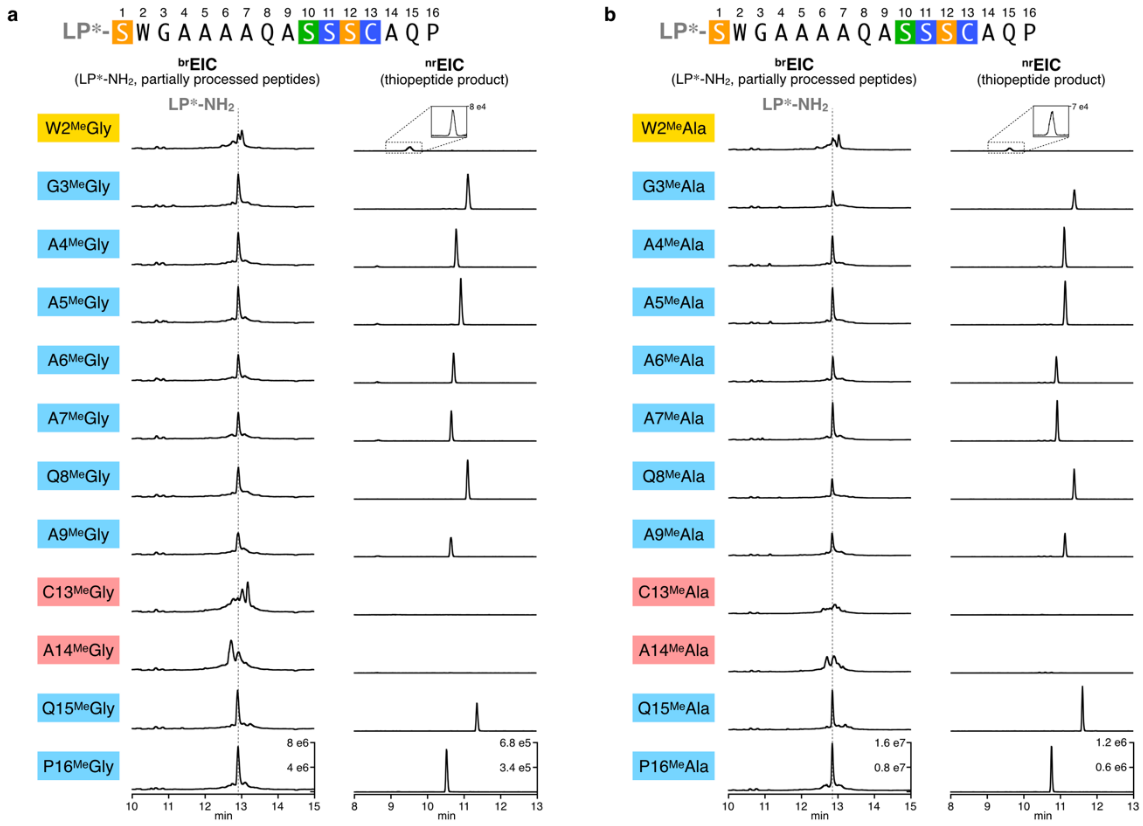

**Figure S18**. Synthesis of *N*-methylated thiopeptides with FIT-Laz. **(a)** $^{Me}$Gly-scanning and **(b)** $^{Me}$Ala-scanning mutagenesis of LazA$^{min}$. Precursor peptides accessed with *in vitro* genetic code reprogramming were treated with the full enzyme set and the reaction outcomes were analyzed by LC-MS. Displayed are LC-MS chromatograms ($^{br}$EIC chromatograms on the left showing partially processed linear peptides and LP$^{*}$-$NH_2$ after enzymatic treatment, and $^{nr}$EIC chromatograms on the right for expected thiopeptides generated at *m/z* 0.10 tolerance window). LP$^{*}$ stands for LazA LP sequence where formyl-Met is replaced with *N*-biotinylated-Phe (see S.I. 2.8 for details).

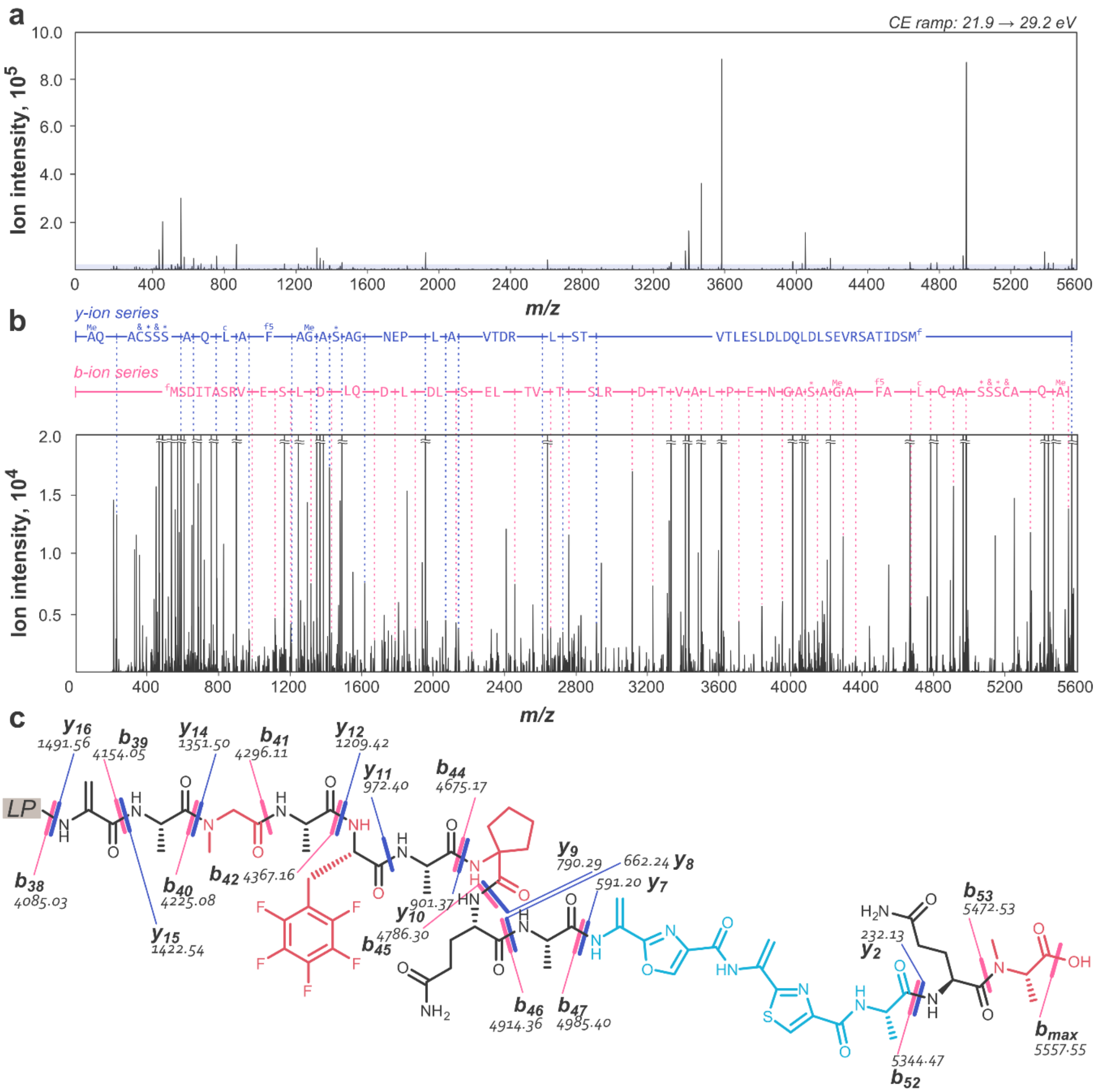

**Figure S19**. MS/MS spectrum of LazA containing 4 npAAs after treatment with LazBDEF/tRNA/GluRS. **(a)** Charge-deconvoluted CID fragmentation spectrum obtained with collision energies ramped from 21.9 to 29.2 eV. **(b)** Y-axis zoom of the shaded area from (a) with spectral assignments; *y*- and *b*-ions are annotated; stable molecule losses ($H_2O$, $NH_3$, CO, etc) and double fragmentation assignments are omitted for clarity; "fM" stands for formyl-methionine; asterisks denote dehydration events; ampersands – azole formations. **(c)** Assigned chemical structure of the LazA CP region with mapped annotations; 2 azoles and 2 dehydrations can be localized to the region highlighted in cyan; positions of npAAs (highlighted in red) and the third dehydration match the displayed structure.

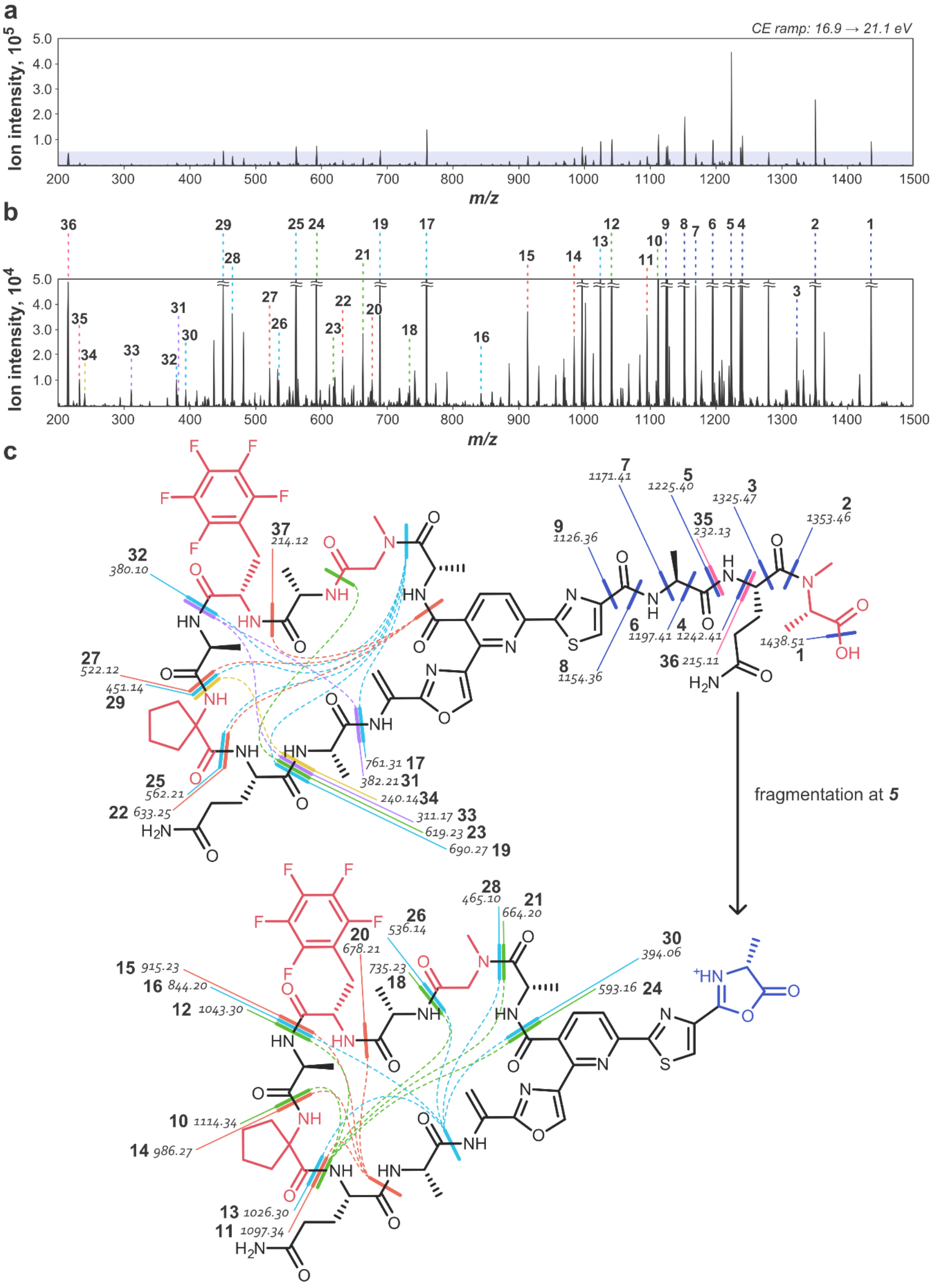

a
CE ramp: 16.9 → 21.1 eV
Ion intensity, 10^5
m/z
b
Ion intensity, 10^4
m/z
c
fragmentation at 5
1 1438.51
2 1353.46
3 1325.47
4 1242.41
5 1225.40
6 1197.41
7 1171.41
8 1154.36
9 1126.36
10 1114.34
11 1097.34
12 1043.30
13 1026.30
14 986.27
15 915.23
16 844.20
17 761.31
18 735.23
19 690.27
20 678.21
21 664.20
22 633.25
23 619.23
24 593.16
25 562.21
26 536.14
27 522.12
28 465.10
29 451.14
30 394.06
31 382.21
32 380.10
33 311.17
34 240.14
35 232.13
36 215.11
37 214.12

**Figure S20**. Annotated fragmentation spectrum of the hybrid lactazole thiopeptide containing 4 npAAs. **(a)** Charge-deconvoluted CID fragmentation spectrum obtained with collision energies ramped from 16.9 to 21.1 eV. **(b)** Y-axis zoom of the shaded area from a) with spectral assignments. **(c)** Assigned chemical structure of the hybrid thiopeptide with mapped assignments. Under the acquisition conditions the thiopeptide underwent multiple double fragmentations allowing the mapping of amino acids within the macrocycle. A prominent *b*-type ion 5 underwent further fragmentations, which are displayed separately. Some ions can have multiple potential assignments. In such cases, only 1 isomer is shown. The spectrum enabled unambiguous assignment of positions of npAAs and post-translational modifications.

### 4. Codon-optimized sequences used for protein production

#### 4.1 LazB codon-optimized ORF nucleotide sequence

CATATGCCGA ATCGCGCCGC ACCCCCTCGT GATGCACGTG CCCCGGTTCC TGCACCGGCC
CCTGCCGCTT ATGCTGCACG TCATGCCCTG GTTCGCAGCA CAGTTCTGGC ATGGCCGGCA
CAGAGCGCAG CCACAGCACA TACCCGTGCC CTGCTGCGCG ATCTGGCAGC CGCAGAAGCA
GCAGCAGAGG CCCTGCGCCC GGCCCTGTGT GATGATCTGT ATGCCGGTCG CGCAGGCCAC
GATGAAGAGT TTCACCGTCG CGTTGTGCTG CCTTTACGCC GCGATCTGCA TAACGGCCGT
ACCCCGCGCG CAGCACTGTT AGACCGCTTA GCAGACCTGC CGCGTCGTAT TCCTCGCCTG
GCCGAATGGC TGGAACTGCG TCGCTTACGC GCACGCCTGC TGGATGCATT AGCCGATGCC
GTTCCGCCTG CACTGGCAGC AGAACGTGCA GCCTTAGCCG ACATTTGCCG TGAACCGGTT
TTCACACGCG CCGTGGCACT GACCAGCGCA GATTTACTGC GCGCAGTTGC CAATACAGCC
GGTGCAACCG GCGAACCGCC TCGCGGCCGT GCCCGTAAAG AAGAAGCAGC CGTTCTGCGC
CACGCACTGC GTGCCACCGC CAAAACCAGC CCTCTGAGCT GGTTCACCGC AGTTGGTTGG
AGTAGCGAGG ATGGCGAAGC AGCAGCCGGT GAACCTCGTG CCTGTGTGCG CGAAAACCGT
GCCCTGGTTA CCGCCCTGGT TCAAGCCCTG CTGGATGACC CTCGTCGCAG TCGTACCCTG
CCGCATCGTA TGACCAGCGC AGCACGCGTG GCAGATGGTC GTGCACGTTA TGCCCGTGCA
GAGGCACTGT TTGCAGGCGG CCGCTTTCTG GTTACCCGCG AAGAAGAAGT GGAACTGGCA
GCCCGCCCTG AGTTAGCACT GCTGGCCAGT TTAGCAGCCA CACCGGCCCC GCCTGACCGT
TTAGCAGCCG GTCTGGCAGA AGCCTTAGGT CGCCCTGGTG GCGATCCTGG CGCACAACGC
TTTGTTGACC AGCTGGTTAC AGCCCGTCTG CTGGTGCCGA CAGAGCCTGT TGATCCTCAG
GATCGTCATC CTCTGCGCAG TCTGGCCGGC TGGTTACGTC AATGGCCGCA AGACGCAGAA
CTGGCACATC GCATTGAGCA GCTGGATCGC CAGAACGCAG AGCTGGCAGT GACAACAGGT
GAACATCGCC CGGAACTGTT AGCCGTGCTG GCCGAGCGTT GGCGTCTGTT ACTGGCCGAT
GCCGGTCGTC CGGTGCCGCA GGAAGCAGCA CCGCTGAGTG TTCTGAGCGA AGACGTTCAC
GCACCGGCCC CTCCGCAACC TCGTCCGGGT GCAGCAGATC GCGCAGCATT AGCCGAACTG
ACAGCCTTAG CCGAGCTGTT TGACCACGCC CATCTGATGC GCCGTGCAGC ACGCGGTCGC
TTTGTGGCAC GCTATGGCGT GGGCGGTGTG TGCGATGCAC CTTGGGATTT TGCCGCCGAC
TTAGCCGACA GTTGGGCAGA TCCGACCCCG CCTGATGAGT TAGCCGCCCT GCGTGAGGAG
TTTGCCAAGC TGCCGGAACA AGATGGTGAG CTGGTTCTGC CGGCAGAACG TATTCGTGCC
CTGGCAGCCC GTTTACCGCA TTGGACAGCC GCACGTCCTT TAAGCTACAG CTGGTTTGTG
CAGCGCGGTA GTGCAGATGG CCTGTTATGC GTGAACCACG TTTACGGCGG TTGGGGTCGC
TTTACCAGCC GTTTTCTGGA CGGTCTGGCA CCTCAGGCCG CAACAGAAGT GGCCCGTCAA
CTGCGCAGTG GTTTAGGTGC CGGTGCACGT GCAGCACAGA TCCGTCCGGT GGGTGGCTTC
AACGCCAACC TGCACCCGCG TCTGTTAGCA GATGAGATTG GTCCTGACCG TCAGTGGACA
GGTCTGGCCG AGAGCGATTT AGACCTGGTT CACGATCCGG TGGACGACCA ACTGCGCCTG
CGCCTGCGTA CAACCGGCGA GCTGCTGGAT GTTCTGTACC TGGGTTTCCT GGCACCGGTG
ATGCTGCCTC GTCGTCTGGG CCCGCTGTTA AATGACCACC CGGAAGGCGT TGTGGATTTC
CGCCCGCTGC TGCCTCGCAC AACCCTGGCC GCACCGGGTG GTACCGTTTT ACGCACACCG
CGCTTACGCC ACCGTCATGT GGTTCTGGCC CGCCGTCGCT GGCTGCTGCC GGCCGGCGTG
TTAGATGCAC TGCGTGCCGA TTTAGCCGCA GATGCAGGTC CTGACGGTGT TCCGGCAGCC
GCAGTGGCAC GTTGGCGCGC ACGTCTGGAC CTGCCTGAAC AACTGTTCTT ACATCCGGCC
CCTGCAGCCG CCGATCCTGC CGGTACCCCG GGCGATGCAT TTGTGGCCCA CCTGCGCGCA
CCGAAACCGC AACCTGTGGA CCTGGGTAAT CCGCTGCACC TGCGCCATCT GGCCAAATGG
CTGACACGCC ATCCTCGTGG CGCCGTTCTG GAAGAGGCAC TGCCGGCAAT CGCCGGTCAT
CCGGAACCTA CCCGCGCCGT GGAACTGGTT GTTGAAACAT ATCGCCCGGG TCGTGGTAGC
GAACAGGCCG CCGGTGCCTT TGAAAGCGTT CGCACCGCAA TTGCAGAAGA AGCCCCTGAC
GAACTCGAG

#### 4.2 LazC codon-optimized ORF nucleotide sequence

```
CATATGAGCG ACCCGGCAGA TGGTCGTGGC GCAGTTACCG CATGGGATGT TGTGCTGTAC
CACTATCGCC CGGATAAAGC CCGTGCACTG CGCGAAGCCG TTCTGCCTCT GGCACGTCAG
GCAGCCGCAG AAGGTCTGGC CGCACACGTG GAACGCCATT GGCGTTTTGG TCCGCATCTG
CGTCTGCGCC TGCGTGGTCC TGAAGCCCGT GTTGCAGGTG CCGCACAGCG TGCAGCAGAA
GCATTACGTG CCTGGGCAGC CGCACATCCG AGTGTTGCCG ATCGCAGTGA CGAACAGCTG
CTGGCAGAGG CCGCAGTTGC AGGTCGCGCA GAGTTAATTG CCCCGCCGTA TGCCCCGCTG
GTGCCGGATA ACACCGTGGT TGCAGCCCCG GCAGATCGTA GCGCAGAAGA TGCCTTACGC
GCCCTGATTG GTGCCGAAAG TGCCGAGCTG CGCGAAGAAC TGCTGCGCAC CGGTCTGCCT
GCCCTGGATA GCGCATGCCA CTTCCTGGGT GCACACGGTG ACACACCGCA GGCCCGTGTG
CAGCTGGTGG TTACCGCACT GGCCGCACAT GCAACCGCAC ACCCTGATGG CCTGGTTGGT
GCCCACTATT CTGTGCTGAG CCATCTGGAA GACTTCCTGG TTCACGAAGA TCCGGATGGT
AGTCTGCGCG CCGCATTTGA ACGCCGTTGG GAACAAAGCG GCCGTGCCGT TACAGCCCTG
GTTGGTCGCA TTGCCGATGG CGGTGCCCGT GATTGGGAAC GTGATTGGGC CCATTGGAGC
GCAACCGCCT GGAGTCTGGC AGAGCGTCGT CTGACAGCCG GTGCAGATCT GGGCGGTCGT
CATGCCGAAT ATCGTGAACG CGCCGAAGCC TTAGGCGATC CGGCCACAGC AGAACGTTGG
AACGCAGAAC TGCGTACCCG CTATAGCGAG TTTCATCGCA TGCTGCAGCG CGCAGACCCG
GATGGTCGTA TGTGGCATCG TCCGGATTAT CTGATTAACC GCGCCGGCAC AAATGGCCTG
TATCGTCTGC TGGCCATCTG CGATGTTCGT CCGATGGAGC GTTATCTGGC AGCCCATCTG
CTGGTGCGCA GTGTTCCTGA GCTGACCGGC CATCGCTGGC AGACCCTGTT AGGCGCCGCA
GAACAGCCGG GTGGTCCTGA ACAGAGTGGT GCAGCAGGCG CAACCGGTGG TGCAGGTCGT
ACCAAACTGG AGGGTGCCGC ACTCGAG
```

#### 4.3 LazD codon-optimized ORF nucleotide sequence

```
CATATGACCG CAGAACCGGA TGCCGTTCGT CCTCGTTTAC GTCCGGGTGT GGCAGTTACA
CCGCTGCGCG AGGGTCTGCA CTTACGCGGT CGTGAAAGCA GCGTGACATT AGAAGGCAGC
CGTGCACTGC CGGCATTATG GCAGGTTCTG GCAGCCCGTT TAGGCCCGCA GGCAGAGGCA
GCAGATGCAG CCGTGGAAGC CACCGTGGAA CCGCGTGTTG CCGCAGCACT GGCAACCGTG
ACAGCACGCC TGCGTGAGCA CGGCCTGCTG GTTGATCACC CGGATGGCGT GCGCTTACCC
CCTTGGCCGG GCGCAGTTGC AGATGATCCG GGCGGTGCCG AGGCAGCATT AGCAGCAGCC
CGTCCGGTTG TTGCCGCAGC AGATCCGGAT GGCCCTAGTG CCCGCGCAAT GGCACGTGCA
TTAGCACGCG GTGGCACCGC AGCACCGGCA GTTGTGGCAG AACCGGGTTT ACCTGCCGGC
CGTGTGGTTG CCACCGCAGA TGGTCCTGCC GGTACCGAAT TAGCAGTGGC AGTGCAGTGC
GGCGCAGACG GTGGCTTTGT TACCGAACCT GCCGACCCGG CACGTGCACG TACAGACGCA
GCAGCATTAG CAGCACGTCT GGAACCTGCA CCGGTTGCCG ACCCGCCGCC GGTTTTATTA
GCACTGCTGG CCGCCGCAGG TGCACAACGC TTACTGTGCG CAGTGGCCGG TCTGCCTGAT
CCTGGTGAAC CTGCAGATGA CCCGCGCTTA CTGGATGGTC GCCCGACCGT GCTGATTGCA
GATGCAGCAC CGCCGCATGC AGAGCATCAT CCTTGGGCCG CAGGCCCTGG TGCAGTTGCA
GCACCTCCGG GCAGTTTAGC CGAAGCCCTG CGCCGTGTGA ATGCCCTGGG TGATCCGCGC
CTGGGCGTGT TAGACGCACC GAGTGCCGGT GACCTGCCGC AGTTACCGGT GGCACTGGTT
AGCTGCGCCA CACCGGCAGG TCCTTTAGCC GCAGGTGCAG TTCGTACCGA CTTAGCACGC
CTGGCAGCAG CATGCCGTAG TGCCGAACTG CATTTAGCAG CAGTGGGTGG TGGCGCAGTG
CCGGTTGTGG GCGTTGATCC GGATCATGCC TTAGGTCTGG CACTGCGTCG CGCCGTTCTG
GCACGCGCAG TTCGTGGTGA CCGTCCTCTG CCGGATGATC GTGCAGCCCG CGGCGATCGT
ACCGTTCCGG AAAGTGCCTG GCGTGAGCAC CCGCAAGCAG GTCATTGGTA CGGTGTGTTA
GCCCGTCGTC TGGGCCGTGC ACCTGAACCT ACCATTCGTC AGCTGAGCGG CGAGAGCGTG
TACCTGGCCC AGGTTGAAGA AGGTCGTGCC GTTGAAGCAA CCCCGGCAGA TGCCGTTGCC
CATGCAGCAC TGGCAGCACT GACCCGCTTA ATGGCCCGTG GTGCAGGCCT GGCAGCAGTG
CATCATACCG TTTTAAGCGG CGCAGCAGCA CCGTTAGCAG CCGCAGGTCG TACCCCTGCA
GCATGGACCG ACCTGGGCTG GGCAGATCGT TGGCTGGCCG ATATTGCCGA TCGTGAAGCC
GACCTGCACG CCGCACTGGT GCGTATTACC GGTTTACGTA CCGCACGTTG GCGTCCGGCA
ACACCTGAAG CCCGTCCGTT CGCAGATGCC CTGGATGGTT GCGGTTTTAC CGCCCTGACA
GCCGAAGGCG GTCGTCCG
```

#### 4.4. LazE codon-optimized ORF nucleotide sequence

```
CATATGAGCG AACTGCCGGT TCTGACACCG GTGGAGGCAC TGGCCGCCAC AAGTGGTACA
GCCGTGGTGC ACCTGACCGA ATGGACCCTG GGTCTGGCAG CCCGCTTAAG CCGCCACGCA
TTAGCACATC CGGTGCGCCT GGTTCCGGTT CGTGAAGACG GCGCCTTAAC CGTGGTGGGT
CCTGTTCTGG CACCTGGCGC ACCTGCATGC CTGGCCTGCG TTGAATATCA GCGTCTGGCA
ACCGCAGGTG GTCGCGTGCC GTGGCAGAGT CCTGCCTTAG CCCTGGGTGG TACCGGTACA
CCGGCATTTG CAGAGGCAGT GACAGCCCTG GCAGCCGAAT TAGCCAAGGG TCCGGAGGCA
GCAGAGAGTG CCGAAGGCGC AGGTAGTCCG GAAGCCGCAG AAAGTGCCGA GGGCGCAGGC
AGTCCTGAAG CCGCCGAAAG CGCAGATGGC GCCGGTAGTC CGGAAGCAGC CGAAAGTGCC
GATGGTGCCG GTAGCCCGGA AGCAGCAGAA AGTGCCGACG GTGCAGGTAG TCCGGAGGCA
GCAGAAAGCG CCGATGGCGC AACAGTTCAT GTTGTGCATG GTGGCCGCGC CACATGGAGC
ACCCATCGTG TTCGCCCTGT TGGCGGCTGC GAGGTTTGTC GCCCGTTACC GCCTGATACC
CCTGAGGCAG CCCGCTTACC TGCCACCCCT CGTCCTCTGC CTGATCCGGC AGTTCTGCGC
GGTCCGAATG ACCGTACAGA TGCCGGCCAG CTGCGTGCAG AACTGTACGA TGAGCGTTTT
GGCCCTGTGC GTCGTCTGTT TCGCACAGAA GATAGCGCCT TTGCACTGAC AACCGCATGG
GTGACAGATG GTCGCGCCCT GGATGATGGC GGCTATGGTC GCGCCGCAGA CTTTCGTAGC
AGCGAACGCG TGGCCCTGTT TGAAGCCGTG GAGCGTCATG CAGGTATGCG CCCTCTGGGC
CGTCGCACAG TGTTACGCGC CAGCTATGCA GAACTGGCCC GTGAGCTGGG CCCGGATGCA
GTTTTAGATC CTGCACGTCT GGGTCTGCCG GATGATCCGC ATCAGGGTCA TCCTACACCG
GCAACAGCCC CGTATACACC TGAGCTGGTG TTAGATTGGG TGCACGGTTG GAGTCTGACC
CGCCGTCGTC CGGTTGCAGT TCCGGAACGT GTTGCCTACT GGGAGGTGCC GGGTCGTGAT
CGTCCTCGTG TGGTGTACGA AAGCAGCAAT GGCTGTGGCC TGGGTAATAG CCCTCAAGAA
GCCGCCCTGT ACGGCCTGTT CGAAGTTGCA GAGCGCGATG CCTTTCTGAT GGCCTGGTAT
GCACGCACAC CGCTGCCTGG CGTTGCCGTT CCTACCGAAG ATCCGCAAAT TGCCGAGTTA
GCAGACCGCG CCGAGTTATT CGGTTATCAC CTGACCCTGC TGGATGCAAC CAACGATCTG
GGTGTGCCGG CCGTGATCGC CCTGTGTCGT CATCGTGGCG ACCATCCGGA CGCACCTCGC
ACATTACTGG CCGCAGGTGC CCACCATGAT CCGCGCACAG CCATTCGTAG CGCCGTTGCC
GAAGTGGTGA CAAATGTTCA AGAAGCTCCT CATCGTAGCA CAGCACCGGG TGGTCCGCGT
GACCCTCAGC GTTTACGTCC GATGCTGGAG CGTCCGGAAC TGGTGGTGAG CCTGGACGAT
CATGTGGGTC TGAACACCCT GCCGGAAGCA CAGCCTCGTC TGGACTTTCT GTTTGCCGGC
CCCCCTCCGG TGCCTTGGAC AGAACGCTGG CCTGGCGATC CGGAACCGGT GACCGATCTG
ACAGATCTGT TAGAGCGCAC AGTGACACGC CTGGCAGGTG AAGATCTGGA GGTGCTGGTT
GTTACCCAGG ATGAGCCGGG CGTTCGTGAC CGCTTAGGTC TGCATTGCGC CAAAGTTGTT
GTGCCGGGTA CACTGCCTAT GACCTTCGGC CATGCAAATC GTCGCACACG CGGCTTAAGC
CGTCTGCTGG AGGTGCCGTA TCGCTTAGGT CGTACACCGG CCCCTTTACG CCACGATGAA
TTACCGCTGC ATCCGCATCC TTTTCCT
```

#### 4.5. LazF ORF nucleotide sequence

```
                           ATGACCA CCCACGCGCT GCCGGCCACC ACCTGGCACA
GCCTGCACCT CGCGCTGCCG CTGCCCGCCC GCGAGGCCGA CGCCTTCCTC ACCGAGGACC
TCGCCCCGCT GATGGACGGG CTCGCCGGCA CCGACTGGTT CTTCATCCGC TACGGCGAGG
GCGGCCCCCA CCTGCGCATC CGCCACCGCG GCCCGGGCCC GGCGCCCGCC TCCCTCGCCG
CCGACCTCAC CCGCCTCGCC ACCCGACGCA CCGCGCCCGA CGGCCCGTTC GCGGACGGGC
ACGGCACCGT GACGGAGGTC CCGTACGAGC CCGAGACCGA GCGCTACGGC GGCGCGGCCC
TGCTGCCGAT CGCCGAAGAG GTGTTCACCC ACTCCACCCG CGCCGCCGTC CGCGCCCTGC
ACGCCCTCGG CGCGGCCCCC GAGAAGCGGT TGCAGCTCGC CCTCGACCTC GCCCACACCA
CCGCGTACGC GCTCGGCCTC GACGAACTCG CCGCCTCCCG CTGGCTGCGC CGCCACGCCG
CCGCCTGGCG CTGGGTCACC GAGTTCCGGC CGCTGCCGGG CGCCGCCGTG CACACCCGGG
TCAACACCGT GTTCGCCCGC CAGCGGGAGA CGCTGGCCCG CCGCGCCCGG GAGCTGCGCG
CGGCACTGGA CGCCGGCACG GCCAGCCCCT GGCTGGGGGA CTGGGCGGCG CGGGCCGCCG
AGGCCGCCGC CCGGATGCGG GCCGTCGCGG CCGCCGAGGC CGCCGCGCCC GCCTCGGCCG
CCGAGGAGGC CGAGGAGCGC CTGGAGTGGA TCTGGGCCTC CCAGCTGCAC ATGCTGTTCA
ACCGGCTCGG CGTCGGCCCC GACGAGGAAC GGGCGGTCTG CCGCCTCGCC GCCCGCACCC
TGCTGGAGAC CGGACAGCCG TTCACCTTCT TCCCCGCCGA CCACCACGCC CCCGACCACC
AGTACCTGGA ACGCAGCAAG TTCCAGATCG GCCGCGGCGA GGACACCGCC CTGCGCGACC
TGCCGCACGA CCCGGCCCCG AGCCCCCGCC CCGGCGACCT CCCGCTGCCC GCCGACCCGC
TGCCGCCCGT CACCCTCGCC GACGCCCTGC GCACCCGCCG CTCCACCCGC GGCCCGCTGA
CCGGCCCGCT CACCGCCGGC GCGCTCGGCG GACTGCTCTG GTCCGCCTTC GCCCCCGCCC
CCGACACCGG CCACCGCCCG TACCCCAGCG CGGGCGCCCT GCACACCGTC CGGCTGCGCC
TGCTCGCCCT CGCCGTCGAC GGCCTGCCCG CCGGCACCTA CCACTGCCTC CCCGAACACC
GCAGCCTGCG CCCGATCGGC CCGGCCCCCG CCCTCGACGA CCTCAAGGCG CTCTCCTCCT
ACCTCTCCCG CCCGGCCGAG GACCCCGACG CCATCGGCGT CGACCGGGCC CCCGCCGTCC
TCGCCGTGTA CCTCGACCTC GCCCGGCTGC GCCGCCGCTA CGGTCTGCGC GCCCTGCGCC
TCGGCGTCCT GGAAGCCGGA CACCTGGCCC AGAACCTGCT CCTGACCTCC GCCGCCTTCG
GCCTCGGCAC CACCCCCCTC GGCGGCCTCC AGGACGACCT CGCCCACGAA CTCCTCGGCC
TGGACGACCT CGGCGAGCCG ATCCAGTACC TGCTCCCGCT CGGCCGGCCG GGGACTGTAC
CGGTGATCAT GGAGTGA
```

#### 4.6. *S. lividans* GluRS codon-optimized ORF nucleotide sequence

```
                                 GTGGCTAGCG CATCCGGCTC CCCCGTACGC
GTCCGTTTCT GTCCGTCCCC CACCGGCAAC CCCCACGTGG GCCTGGTCCG CACCGCCCTG
TTCAACTGGG CCTTCGCGCG CCACCACCAG GGCACCCTGG TCTTCCGCAT CGAGGACACC
GACGCCGCCC GCGACTCCGA GGAGTCGTAC GACCAGCTGC TCGACTCGAT GCGCTGGCTG
GGCTTCGACT GGGACGAGGG TCCCGAGGTC GGCGGCCCGC ACGCGCCGTA CCGCCAGTCG
CAGCGCATGG ACATCTACCA GGACGTCGCC CAGAAGCTCC TGGACGCCGG CCACGCCTAC
CGCTGCTACT GCTCCCAGGA GGAGCTGGAC ACCCGCCGCG AGGCCGCCCG CGCCGCCGGG
AAGCCCTCCG GCTACGACGG CCACTGCCGC GAGCTGACCG ACGCACAGGT CGAGGAGTAC
ACGTCCCAGG GCCGCGAGCC CATCGTCCGC TTCCGGATGC CCGACGAGGC GATCACCTTC
ACGGACCTGG TCCGCGGCGA GATCACCTAC CTGCCGGAGA ACGTCCCGGA CTACGGCATC
GTCCGCGCCA ACGGGGCGCC CCTCTACACG CTGGTCAACC CCGTCGACGA CGCGCTGATG
GAGATCACCC ACGTCCTGCG CGGCGAGGAC CTGCTCTCCT CCACCCCGCG CCAGATCGCC
CTGTACAAGG CGCTGATCGA GCTGGGCGTC GCCAAGGAGA TCCCCGCCTT CGGCCACCTG
CCGTACGTCA TGGGCGAGGG CAACAAGAAG CTCTCCAAGC GCGACCCGCA GTCGAGCCTC
AACCTCTACC GCGAGCGCGG CTTCCTCCCC GAGGGCCTGC TCAACTACCT CTCCCTCCTC
GGCTGGTCGC TCTCGGCCGA CCAGGACATC TTCACGATCG AGGAGATGGT CGCGGCCTTC
GACGTCTCCG ACGTCCAGCC CAACCCGGCC CGCTTCGACC TCAAGAAGTG CGAGGCGATC
AACGGCGACC ACATCCGCCT GCTGGAGGTC AAGGACTTCA CCGAGCGCTG CCGCCCCTGG
CTGAAGGCCC CCGTCGCCCC CTGGGCGCCG GAGGACTTCG ACGAGGCCAA GTGGCAGGCG
ATCGCGCCGC ACGCGCAGAC CCGCCTGAAG GTCCTCTCCG AGATCACCGA CAACGTCGAC
TTCCTGTTCC TGCCGGAGCC GGTCTTCGAC GAGGCCAGCT GGACCAAGGC CATGAAGGAG
GGCTCGGACG CGCTCCTGAC CACGGCCCGC GAGAAGCTGG ACGCCGCCGA CTGGACCTCC
CCGGAGGCCC TCAAGGAGGC CGTCCTGGCC GCCGGTGAGG CCCACGGTCT CAAGCTCGGC
AAGGCCCAGG CCCCCGTCCG CGTCGCCGTC ACCGGCCGCA CGGTCGGCCT GCCCCTCTTC
GAGTCCCTGG AGGTCCTGGG CAAGGAGAAG GCACTGGCGC GCATCGACGC GGCGCTGGCG
CGACTGGCGG CGTAA
```
